# A genetic screen unveils key molecular steps in the polymerization cycle of the bacterial actin-like MreB

**DOI:** 10.64898/2026.07.23.740262

**Authors:** Alba de San Eustaquio-Campillo, Charlene Cornilleau, Sana Afensiss, Catalina Marchioni, Hind Oulkfif, Lilia Huynh, Davy Martin, Lars Renner, Rut Carballido-López, Louis Renault, Arnaud Chastanet

**Affiliations:** Micalis Institute, INRAE, AgroParisTech, Université Paris-Saclay, 78350 Jouy-en-Josas, France; Université Paris-Saclay, INRAE, UVSQ, VIM, 78350 Jouy-en-Josas, France; Leibniz Institute of Polymer Research, and the Max-Bergmann-Center of Biomaterials, Dresden, Germany; Université Paris-Saclay, CEA, CNRS, Institute for Integrative Biology of the Cell (I2BC), Gif-sur-Yvette, France

**Keywords:** actin-like, MreB, cytoskeleton, ATPase, polymer, lipids

## Abstract

MreB, a bacterial actin homolog and polymerizing ATPase, is central to cell-shape maintenance and cell-wall integrity. Its functions rely on its ability to assemble into dynamic, membrane-associated polymers. However, how nucleotide binding and hydrolysis, MreB-MreB contacts, and membrane association are coordinated to enable polymer assembly and disassembly remains unclear. Here, we combined genetics and live-cell microscopy with biochemical approaches to dissect these processes. Using a highly sensitive reporter of MreB activity, we identified, through a genetic screen, residues critical for MreB function in *Bacillus subtilis*. Subsequent extensive characterization of corresponding stable variants of the homologous *Geobacillus stearothermophilus* MreB revealed that ATP binding, but not ATP hydrolysis, is required for polymerization. Productive longitudinal intraprotofilament contacts are required for efficient ATP hydrolysis and enhance membrane association. Perturbations predicted to weaken lateral interprotofilament contacts altered membrane association and modulated ATPase activity. Together, these effects provide experimental evidence consistent with long-range functional coupling among the longitudinal and lateral protofilament interfaces, the distant nucleotide-binding site, and membrane association dynamics. Moreover, impaired ATP hydrolysis delays disassembly of lipid-associated polymers, indicating that hydrolysis promotes polymer turnover. These results establish key mechanistic steps coordinating ATP-driven MreB polymerization and turnover and provide a basis for a complete MreB assembly–disassembly cycle and for further elucidating how MreB dynamics contribute to cell-wall organization.

## Introduction

Cytoskeletal proteins play a fundamental role in both eukaryotic and bacterial cells, contributing to processes such as cytokinesis, chromosome partitioning, and cell shape maintenance. In bacteria, the essential protein MreB, a homolog of eukaryotic actin, preserves cell shape by controlling the synthesis of the cell wall (CW) ^1, 2^. The CW forms a stiff extracellular shell that confers physical integrity to most bacterial cells. Also known as the sacculus, it is a complex structure mainly composed of a continuous, load-bearing peptidoglycan (PG) network enclosing the cytoplasmic membrane. Disruptions or defects in the PG synthesis result in cell deformation, swelling, and ultimately lysis, explaining why many clinically important antibiotics target this process. Consequently, most proteins involved in PG synthesis are essential or otherwise critical for bacterial survival.

The current consensus supports the existence of PG synthesis machineries, with the “Rod-complex” being the primary driver of longitudinal PG expansion during growth ^3^. Although its precise composition and status as a holoenzyme remain uncertain, a core set of key proteins has been proposed. This set includes the proteins carrying the enzymatic activities responsible for PG extension (RodA, PbpH, PBP2A) which synthesize the PG strands and link them to the existing meshwork, parallel to one another and perpendicularly to the cell long axis. The complex would also contain regulatory proteins, including MreC, RagB, and RodZ, and the essential accessory protein MreD, whose precise function remains unclear ^3, 4^. Among these proteins, MreB is a central component: its loss does not abolish but disorganize PG synthesis, rapid loss of cell shape and CW integrity, and ultimately lysis.

MreB is a cytoskeletal protein and a homologue of eukaryotic actin, sharing its three-dimensional fold and its ability to polymerize ^5^. The protein is encoded by a single gene in most phyla, but paralogs are frequent in Gram-positive (G(+)) *Bacilli* (*e.g.*, the *Bacillus subtilis* genome encodes MreB, Mbl, and MreBH ^2^), while wall-less *Spiroplasma* (Mollicutes, which evolved from G(+) ancestors) carry 5 to 8 genes ^6, 7^. Genetically encoded fluorescent labelling revealed that MreB assembles into distinctive patterns of membrane-associated, dynamic, discrete foci *in vivo*, which represent short filaments below the diffraction limit of light microscopy ^1, 8^. The processive motion of these foci around the cell circumference, along their filament axis, depends on active PG synthesis ^8, 9, 10^. Thus, MreB’s directional motion is considered to reflect the motion of the Rod complex driven by PG synthesis. The ability of MreB polymers to deform liposomes *in vitro* suggested that membrane-bound MreB filaments align with the highest negative curvature, hence circumferentially along the cell’s short axis ^11^. Therefore, MreB’s main function may be to guide the PG elongation complex around the cell circumference, ensuring organized synthesis.

MreB structures from both G(+) and G(-) bacteria have been solved by X-ray crystallography, revealing an actin-like fold, with 4 subdomains (IA, IB, IIA, IIB) surrounding a central catalytic cleft that binds and hydrolyzes ATP or GTP ^5, 12, 13, 14, 15^. However, whereas most actin-family members assemble into twisted helices, MreB forms flat double-stranded polymers ^16^, probably with an antiparallel organization ^13, 17^. Another distinctive property of MreB is its direct association with lipids through hydrophobic and/or positively charged residues ^12, 14, 18^. This interaction is essential for MreB localization and function *in vivo* ^18^, and is required for efficient polymerization of G(+) MreB ^12^. Hydrophobic membrane-binding regions have long complicated the purification and study of MreB from the classical G(-) and G(+) model organisms, *E. coli* and *B. subtilis*, respectively ^12, 19^. Recently, we used the more soluble and stable MreB from *Geobacillus stearothermophilus*, a thermophilic G(+) bacterium closely related to *B. subtilis,* to determine the first G(+) MreB structure and identify features required for G(+) MreB membrane association, as well as a dual requirement for ATP and membrane binding in polymer formation ^12^. However, the molecular determinants governing MreB assembly and disassembly, membrane association and dissociation, and the respective roles of ATP binding and hydrolysis in these processes remain poorly understood. It is also unclear how these biochemical properties influence MreB dynamics in cells.

Although numerous MreB mutants have been generated over the past three decades, most were selected as gain-of-function mutations conferring resistance to chemical or protein inhibitors targeting MreB ^20, 21, 22, 23, 24, 25^. A notable exception is the recent alanine-scanning mutagenesis of *E. coli mreB* ^26^. Moreover, only a handful of mutants have been studied in Firmicutes, *i.e.* walled G(+) bacteria ^10, 20, 27^. We therefore developed a mutagenesis screen in *B. subtilis* to isolate loss-of-function mutants and identify residues critical for MreB functions. Using a functional *mreB-gfp* fusion expressed at wild-type levels, we identified mutants defective in cell morphology and MreB localization in vivo. We then combined genetic analysis and live-cell microscopy with biochemical and biophysical characterization of purified *G. stearothermophilus* MreB variants. This complementary in vivo and in vitro analysis shows that ATP binding, but not hydrolysis, is required for polymerization, whereas ATP hydrolysis promotes polymer disassembly. Partial biochemical deficiencies can also translate into perturbed dynamics *in vivo*. Finally, productive MreB–MreB interactions are required for efficient ATP hydrolysis and membrane adhesion, while preventing them suppresses futile ATP hydrolysis. Together, these findings identify key mechanistic steps linking nucleotide binding and hydrolysis, MreB–MreB contacts, membrane association, and polymer assembly and disassembly, providing a basis for an ATP-driven MreB assembly–disassembly cycle.

## Results

### A genetic screen reveals mutations affecting MreB function in *B. subtilis*

We aimed to design a simple, sensitive screen to identify mutants with impaired MreB function. We previously reported that, while MreBH (one of the two paralogs of MreB) is barely detectable in a wild-type strain, it accumulates to high levels in a mutant deleted for *mreB* ^28^. We reasoned that induction of the *mreBH* promoter in the absence of functional MreB could provide a suitable reporter. To test this idea, we constructed a transcriptional reporter fusion between the promoter region of *mreBH* and *lacZ* (see the Methods section). In an *mreB* knock-out mutant, the reporter showed a strong induction, whereas no expression was detected when placed in a wild-type background (Fig. Sup 1A). Using this fusion as a reporter for MreB functionality, we performed random mutagenesis of the *mreB* locus by low-fidelity PCR to generate mutated fragments, transformed the resulting products into the recipient *B. subtilis* strain carrying the reporter, and screened for blue colonies (see Materials and Methods for details; Fig. 1; Fig. Sup 1B). Out of the 73 mutants selected, approximately half carried a premature stop codon, a frameshift or an extragenic mutation, and one quarter displayed more than one intra-or extragenic single nucleotide polymorphism (SNP) (Fig. Sup 1C), for a total of 51 unique mutations identified. To eliminate any potential additional mutations outside the sequenced *mreB* locus and enable analysis of mutant-protein localization by fluorescence microscopy, we constructed *gfp-mreB* translational fusions, each carrying a single mutation, into a fresh background allowing us to backcross and, when required, isolate the mutations (Fig 1; Fig. Sup. 1D), as described in the Materials and Methods section. For this purpose, we used as a template a *gfp-mreB* fusion inserted at its native locus, under the control of its natural promoter and optimized to express wild-type levels of MreB (strain RCL421; see Materials and Methods section for details). Ultimately, 33 mutants were successfully constructed, each strain carrying a single mutation in *mreB* identified during the screening step (Fig. Sup 1C). The mutations were distributed throughout the coding sequence, spanning roughly 85% of the open reading frame (Fig. Sup 1E).

**Figure 1.**
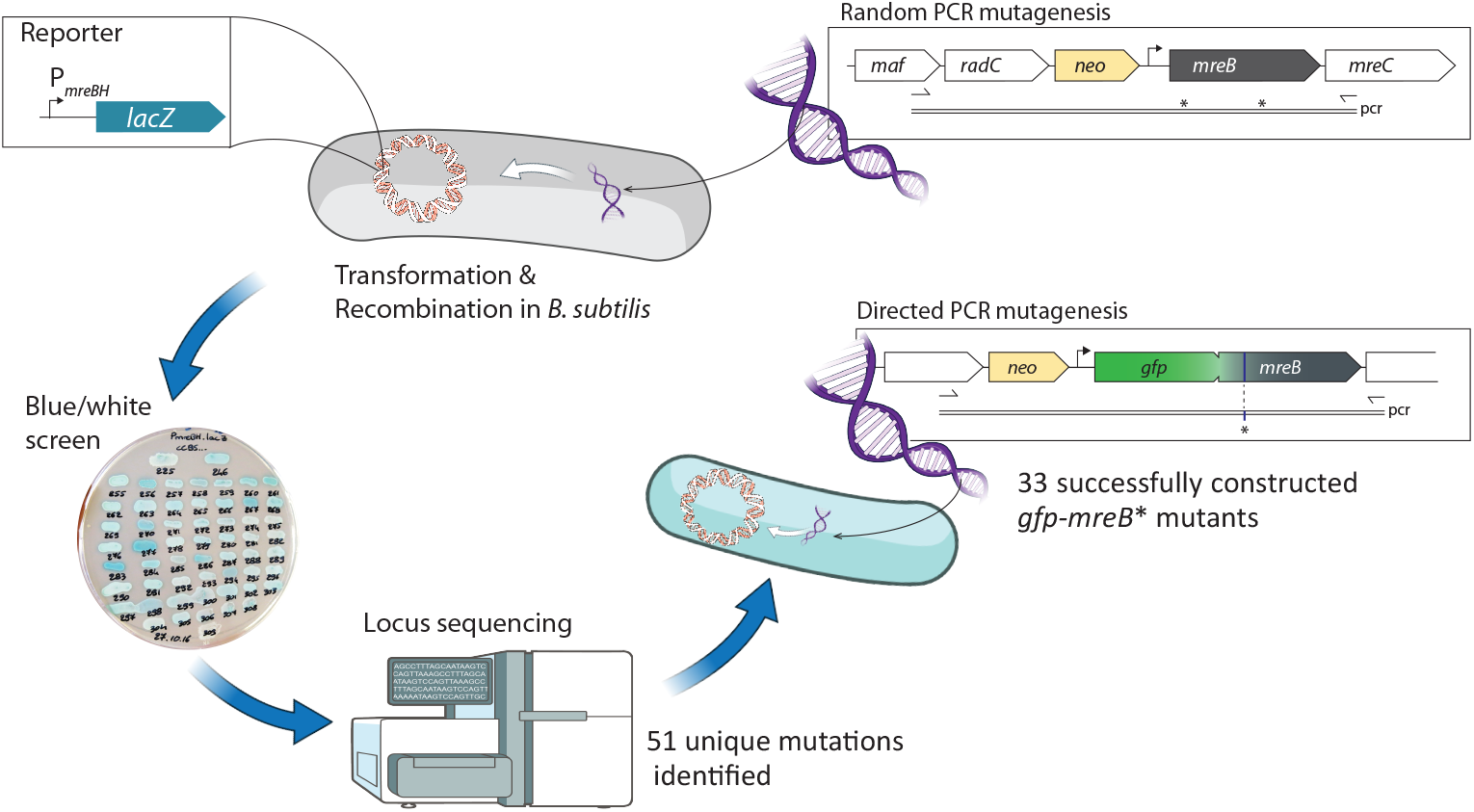
Schematic of the screen designed to identify critical residues for MreB function in *B. subtilis*. A low fidelity polymerase chain reaction is used to amplify the *mreB* locus from a strain carrying a neomycin resistance cassette (RCL414); the resulting product is transformed into naturally competent B. *subtilis* recipient strain carrying the reporter gene (RCL422), and resistant transformants are screened for blue colonies. *mreB* locus of the selected candidates are sequenced to identify the mutations, and new strains were generated carrying a *gfp-mreB* fusions with a single mutation, at the natural locus. Illustrations (bacteria, arrows, sequencer) from NIAID NIH BioArt Source (bioart.niaid.nih.gov/bioart).

### A selection of mutants displaying a complete loss of function

In the present study, we focused on mutants with severe loss of MreB function. To this end, we sub-screened our MreB-deficient strain collection by microscopy for mutants with the most dramatic phenotypes. By comparing the mutants with their wild-type parent (*gfp-mreB^WT^*; RCL424) and the null Δ*mreB* mutant (RCL423), we selected the 5 strains displaying the strongest shape defects during exponential growth in rich LB medium (Fig. 2A). The mutated residues mapped either to the nucleotide-binding/catalytic site (Fig. 2B, blue; G^14^E, G^160^R) or to the longitudinal, intraprotofilament interface, between consecutive monomers (Fig. 2B, orange; G^56^R, L^171^P, G^231^D). These assignments derive from analysis of protein–nucleotide and protein–protein contacts in two crystal structures containing protofilament arrangements (Fig. Sup. 1E; Sup. Mat.). As shown in Fig. 2A, cells carrying the mutated versions of MreB were strongly misshapen and swollen, phenocopying the *mreB* knock-out (Fig. 2A.; Fig. Sup 2A). Because *mreB* inactivation is associated with increased cell width, we used this parameter to quantitatively compare the effects of the mutations (Fig. 2C). All mutants displayed a greater cell width than the wild type, comparable to that of the *mreB*-inactivated strain, except G^56^R, which showed an intermediate average width. Western blotting using anti-MreB antibodies revealed that G^231^D reduced MreB levels, suggesting protein destabilization, whereas the other mutations had limited or no impact (Fig. 2D, Sup 2C). This reduction is unlikely to account for the shape defect, since we previously showed that a strain carrying a non-optimized P*_nat_ gfp-mreB* construct, which drives to similarly low expression levels, displayed an almost wild-type morphology ^29^. We next compared the localization of the fluorescently tagged MreB mutant proteins with that of wild-type MreB by epifluorescence microscopy. The wild-type protein displayed its characteristic distribution as discrete foci along the cytoplasmic membrane ^30^, consistent with the formation of short membrane-associated polymers (Fig 2A). By contrast, all 5 mutant proteins appeared uniformly distributed throughout the cytosol, consistent with a loss of function and severe morphological defects (Fig. 2A.; Fig. Sup 2A). Thus, these mutations prevent the formation of detectable membrane-associated MreB assemblies *in vivo.* Because localization alone cannot distinguish impaired membrane association from defective membrane-dependent polymer assembly, we therefore investigated how these mutations affect the biochemical steps underlying MreB polymerization and membrane association.

**Figure 2.**
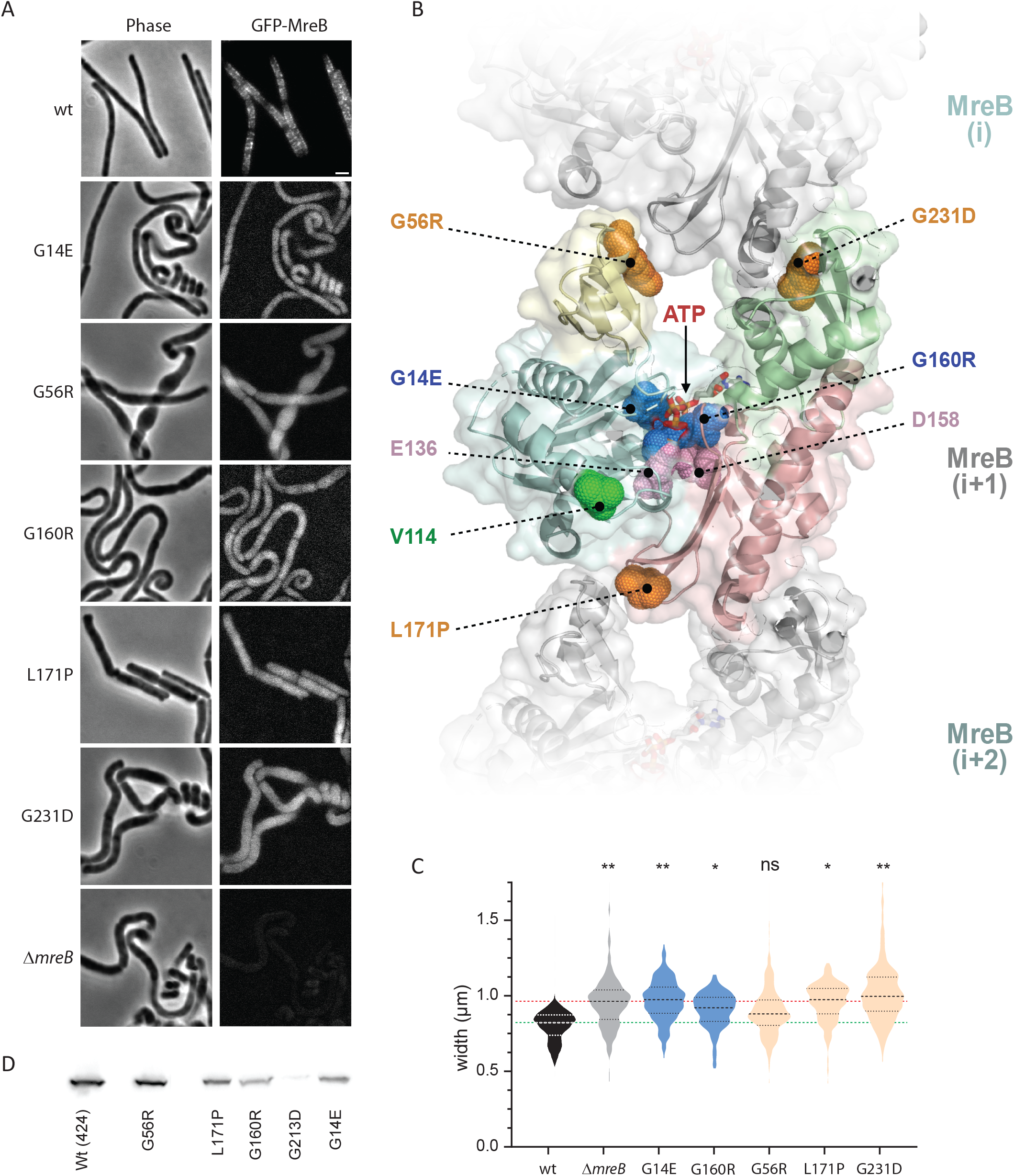
*In vivo* phenotypes of *mreB* point mutants phenocopying a Δ*mreB* KO strain. **A.** Microscopy images of strains carrying a wild type (wt; RCL424) or mutated *gfp-mreB* fusion, compared to a stain deleted for *mreB* (Δ*mreB*; RCL423), observed in phase contrast (Phase) and epifluorescence (GFP-MreB), during rapid exponential growth in rich medium. Scale bar, 1 µm. **B**. Localization of the analyzed mutations in the *G. stearothermophilus* MreB-ATP structure, (PDB 7ZPU; Mao *et al*., 2021). The image includes MreB–MreB contacts generated by crystal packing that may resemble longitudinal intraprotofilament interfaces. The five residues whose mutation causes severe morphological defects are highlighted as enlarged side-chain dots, together with V114, E136 and D158. Residues are color-coded according to their predicted functional roles: intraprotofilament contacts: G^56^R, L^171^P and G^231^D (orange); catalytic site: G^14^E and G^160^R (blue), E^136^A and D^158^A (pink); inter-protofilament contact: V114 (green; highlighted in Fig. Sup. 2G). The four subdomains of the central MreB subunit, shown as ribbons, are distinguished by color: IA (pale blue), IB (pale yellow), IIA (pale red), and IIB (pale green). **C.** Comparative cell width of the strains, determined on cells in A. Data are a compilation of at least two independent experiments. Statistical significance between wild type and mutant’s width was calculated using the Welch’s non-parametric unpaired t-test (**= 0.001<*P*<0.01; *= 0.01<*P*<0.05; ns = *P*>0.05). **D**. MreB levels of the same strains as in A, compared by western blot.

### MreB mutants defective in localization *in vivo* are impaired in polymerization *in vitro*

To further study the effect of the mutations, we purified the mutated proteins and characterized them thoroughly *in vitro*. For this purpose, we turned to MreB from the G(+) thermophilic bacterium *G. stearothermophilus*. We recently showed that MreB from this organism shares a high degree of sequence identity with its ortholog in *B. subtilis*, while exhibiting much better solubility ^12^. Importantly, the residues of interest are conserved between the two species, as are most of the selected mutations (Fig. Sup. 1E). Using site-directed mutagenesis of an expression vector carrying the wild-type *mreB* gene from *G. stearothermophilus* (pCC110; ^12^) (see M&M), we cloned the five aforementioned mutated versions of the gene into an *E. coli* heterologous expression host and purified the corresponding recombinant proteins. All five proteins were purified to homogeneity following our previously established protocol ^12^ (Fig. Sup 3A).

A first analysis by circular dichroism (CD) spectroscopy revealed altered spectra for the G^231^D and L^171^P mutants compared with the wild-type protein (Fig. 3A). This suggests that either the kink introduced by the proline at position 171 or the aspartate side chain at position 231 causes severe structural perturbations within MreB subdomains 3 (IIA) and 4 (IIB), respectively (Fig. 2B), potentially destabilizing the intrinsic MreB fold and accounting for the loss of function of these mutants. Because of these substantial structural alterations, we excluded these two mutants from further analysis. In contrast, all other mutants displayed wild-type CD spectra, suggesting that their overall structure remained unaffected; these mutants were therefore further characterized. We next tested their ability to form polymers *in vitro*. We previously demonstrated by transmission electron microscopy (TEM) that purified wild-type *G. stearothermophilus* MreB forms polymers in the presence of ATP and lipids ^12^. We also developed a semiquantitative TEM assay (sqTEM) to compare the polymerization efficiency across conditions ^12^. Using this methodology, the mutants were incubated under polymerizing conditions, *i.e.*, in the presence of ATP and lipids (see the Methods section for details), and assayed by sqTEM. The G^14^E, G^56^R and G^160^R mutants showed severely reduced or undetectable polymerization under our conditions (Fig 3B), consistent with the absence of detectable foci *in vivo* (Fig. 2A.; Fig. Sup 2A; Sup. Table 4). Because these mutations affect distinct structural regions, we next investigated which biochemical steps in ATP-dependent MreB assembly were disrupted.

**Figure 3.**
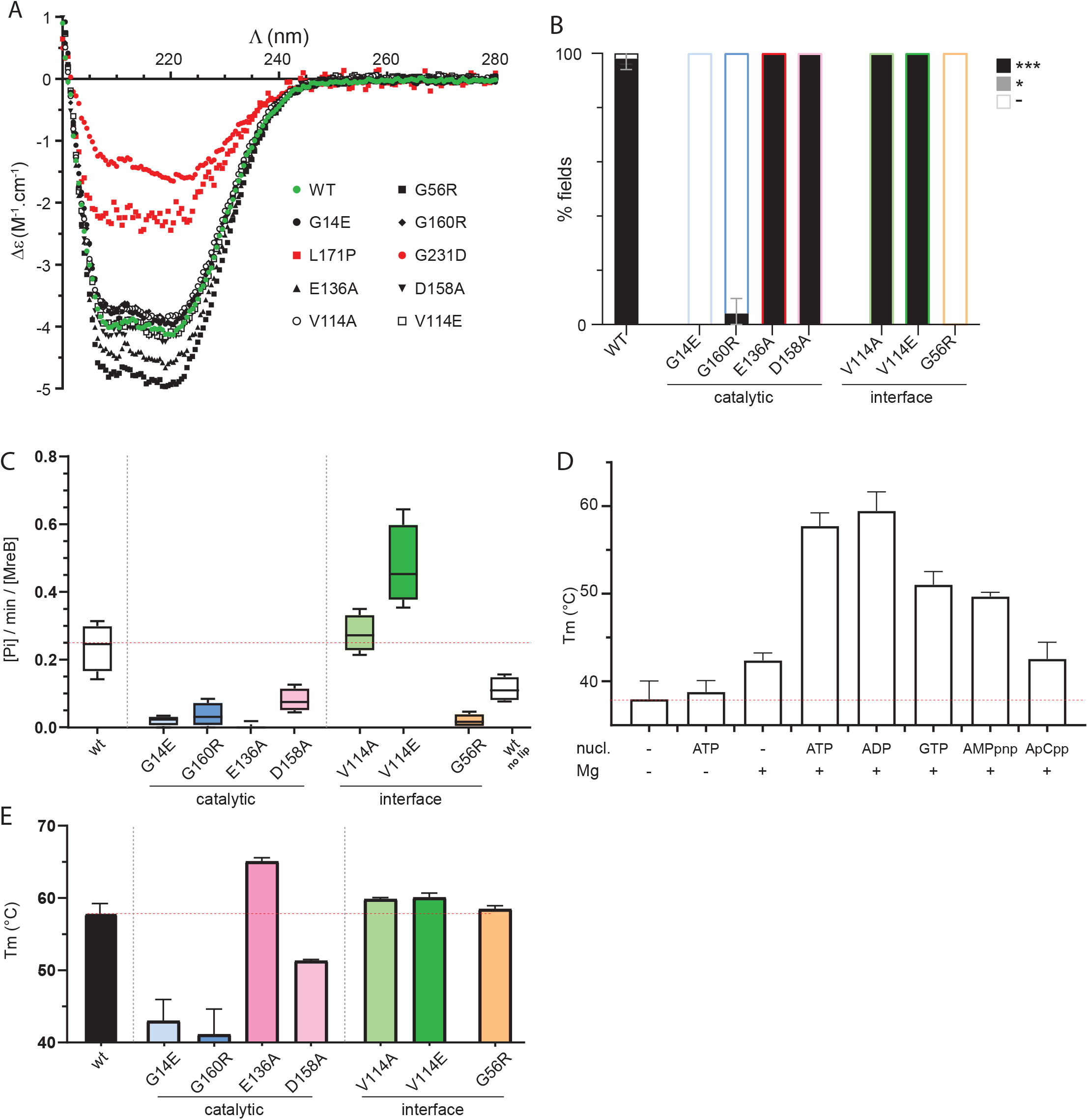
*In vitro* phenotypic characterization of MreB point mutants. **A.** Circular dichroism (CD) spectra of purified wild type (WT; green dots) and mutant MreB proteins, showing similar folding of all mutants except L^171^P (red squares) and G^231^D (red dots). Displayed are typical spectra obtained by averaging four consecutive scans. **B.** Mutants G^14^E, G^56^R and G^160^R are defective for polymer formation. sqTEM analysis, showing the frequency and density of polymer formation observed on negative stained EM images, of purified wild type and mutants of MreB, set to polymerize in standard conditions. **C.** All mutants are defective for hydrolytic activity except V^114^A and V^114^E. Specific activities were determined from the release of inorganic phosphate (Pi) after 2h in the presence of a range of protein concentrations. All experiments were done in standard polymerization conditions except the control without lipids (right hand plot). **D.** Thermal stabilization of wild-type MreB induced by nucleotides and Mg^2+^ binding. Relative nucleotide-induced thermal shifts were assessed by thermal shift assay (TSA), in which the binding of a ligand increases the resistance to thermal denaturation and induces a shift of the melting temperature (Tm). Tm were determined in the presence of 2.5 µM of wild type MreB, 50 µM of ligands (ATP, ADP or GTP), and in the presence or absence of 2 mM Mg^2+^. **E.** Compared ATP-induced thermal stabilization of wild type (WT) and mutant of MreB, determined by TSA. Melting temperature was recorded in the presence of 2.5 µM of protein, 50 µM ATP and 2 mM Mg^2+^. B to E: values are averages of at least two independent experiments and error bars are standard deviations to the mean.

### ATP binding but not hydrolysis is required for MreB polymerization

We aimed to understand how each mutation affected MreB properties and, in turn, infer from these defects how polymerization occurs. We first focused on the catalytic core of MreB and two mutants from our screen affecting its catalytic cleft: G^14^E and G^160^R. We previously showed that *G. stearothermophilus* MreB hydrolyses ATP *in vitro* ^12^. Based on its three-dimensional structure ^12^, the side chains introduced by these mutations are predicted to lie in the ATP binding pocket, clash with the phosphate moiety of the nucleotide, and thereby alter nucleotide binding and hydrolysis (Fig. 2B; Sup. Mat. C). To strengthen our study, we added two additional mutants, D^158^A and E^136^A, predicted to be defective in ATP hydrolysis ^10, 27^, the latter based on a homologous mutation in the related ParM protein ^31^. Both mutations were previously studied *in vivo* in *B. subtilis* but not biochemically characterized *in vitro. In vivo,* they were shown to cause minor morphological defects and limited alterations in MreB localization ^10^. In the wall-less bacterium *S. citri,* the corresponding purified mutants (D^156^A and E^134^A, respectively) showed moderate and strong defects in ATP hydrolysis, respectively ^14^. The D^158^A and E^136^A mutants from *G. stearothermophilus* were produced in an *E. coli* host, purified to homogeneity, and showed broadly preserved secondary structure by CD (Fig. 3A). We assessed the ATPase activity of the four mutants by measuring the release of inorganic phosphate (P_i_) using a malachite green colorimetric assay. All mutants showed reduced specific ATPase activity, with a 3- (D^158^A), 6- (G^160^R), 12- (G^14^E), and 55-fold (E^136^A) reductions relative to wild-type MreB (Fig 3C; Fig Sup 3B).

We next compared the mutants’ ability to bind ATP. We recently estimated the binding kinetics of *G. stearothermophilus* wild-type MreB toward ATP and ADP using the fluorescent nucleotide analog N^6^-(6-amino)hexyl-ATP/ADP-ATTO-488 and fluorescence anisotropy ^32^. Here, we first used isothermal titration calorimetry (ITC) with native ATP and ADP nucleotides. Both nucleotides bound MreB with high affinity in the presence of Mg²⁺, with dissociation constants in similar nanomolar range. ADP bound approximately twofold more tightly than ATP (K_D_ = 42 ± 11.2 nM versus 84 ± 12.1nM, respectively) (Fig Sup 3C). This nucleotide preference differs from that of its eukaryotic homologue G-actin, which binds ATP more tightly than ADP. In the presence of Mg²⁺, ATP binds G-actin with an affinity of approximately 1.2 nM, about 70-fold higher than for MreB, and approximately fourfold more tightly than ADP ^33, 34^. The relatively similar affinities of MreB for ATP and ADP, together with its slight preference for ADP, suggest that nucleotide occupancy may respond sensitively to physiological fluctuations in the intracellular ATP/ADP ratio, which is approximately 2 *in B. subtilis* ^35^. Despite their similar affinities, ATP and ADP displayed markedly different thermodynamic signatures. ATP binding was almost entirely enthalpy-driven (ΔH ≈ −9.56 kcal·mol⁻¹; −TΔS ≈ −0.10 kcal·mol⁻¹), whereas ADP binding combined a substantially smaller favorable enthalpy with a much larger favorable entropic contribution (ΔH ≈ −4.36 kcal·mol⁻¹; −TΔS ≈ −5.7 kcal·mol⁻¹). These results indicate that ATP and ADP achieve similar overall binding free energies through distinct energetic mechanisms.

To compare nucleotide binding across a wide range of conditions and MreB variants, we next turned to the thermal shift assay (TSA), in which ligand binding stabilizes a protein against thermal denaturation, leading to an increased melting temperature (T_m_, defined as the temperature at which half the population is unfolded). For wild-type MreB, T_m_ increased from 38.0 ± 2.1 °C for apo MreB to 42.4 ± 0.9 °C in the presence of Mg^2+^, consistent with cation binding (Fig. 3D). ATP did not affect T_m_ in the absence of Mg^2+^, whereas ATP and Mg^2+^ together increased it to 57.7 ± 1.5 °C, indicating the effective Mg²⁺-dependent nucleotide binding (Fig. 3D). ADP produced a similar increase in T_m_, supporting our ITC observations. GTP, previously shown to support MreB polymerization *in vitro* ^12^, produced a slightly smaller thermal shift than ATP or ADP, similar to that of the non-hydrolysable ATP analog AMPpnp, whereas ApCpp (another non-hydrolysable analog) produced the smallest shift. Our previous failure to observe MreB polymers in the presence of AMPpnp, despite its efficient binding, suggests that this ATP analog does not properly mimic ATP when bound to the active site ^12^. It may induce a different conformation affecting MreB properties such as the critical concentration for polymerization. We next compared the binding of ATP by the 4 mutants affected in their catalytic site in the presence of excess Mg^2+^ (Fig. 3E, Fig. Sup 3D). As predicted from the structure, G^14^E and G^160^R retained virtually no nucleotide-binding capacity, whereas E^136^A and D^158^A did. Interestingly, E^136^A showed the highest T_m_ (65.5 ± 0.5 °C) of all mutants, which, combined with its extremely low ATP hydrolysis rate, suggests that E^136^A traps ATP in a highly stable complex. By contrast, ATP-bound D^158^A displayed a much lower T_m_ than wild-type MreB, indicating reduced thermal stabilization by ATP, which could partially account for its moderately decreased hydrolysis rate (Fig. 3C). When assessed for polymer formation *in vitro* by sqTEM, E^136^A and D^158^A formed polymers indistinguishable from those of wild-type MreB, despite their reduced ATP hydrolysis rate, in a striking difference with G^14^E and G^160^R (Fig. 3B; Fig. Sup 3E). Thus, ATP binding, but not efficient ATP hydrolysis, is required for MreB polymerization (Sup. Table 4). This distinction prompted us to examine next how productive MreB–MreB interactions couple polymer assembly to ATP hydrolysis.

### Productive longitudinal MreB–MreB interactions promote ATP hydrolysis

We next investigated mutations affecting predicted MreB-MreB interfaces. In addition to G^56^R, we examined another mutant from our screen carrying V^114^A, despite its very mild phenotype (Fig. 2B, green; Fig. Sup 2A, 2H). In a previous study, mutation of the homologous residue in *Caulobacter crescentus* MreB (V^118^E) was shown to disrupt lateral interactions between protofilaments (or inter-protofilament contacts) ^13^, making this residue particularly relevant to our study (Fig. sup. 1E; Sup. Mat. A, B). Because the *C. crescentus* mutant carried a glutamate rather than the alanine substitution in our mutant, we also constructed a second *B. subtilis* strain carrying V^114^E. The two mutants presented only limited morphological defects, with rare abnormal flattened poles and twisted cells, features frequently observed in *mreB* knock-outs (Fig. Sup 2A). Although expression of the *lacZ* reporter was low in mutant V^114^A, explaining why it was not initially selected, V^114^E showed strong induction of the *mreBH* promoter, a response typical of impaired MreB function (Fig. Sup 2G). Consistent with a stronger effect of the glutamate substitution, V^114^A cells had a more wild-type-like width, whereas V^114^E cells were slightly enlarged (Fig. Sup 2B). The localization of both GFP-MreB mutants appeared indistinguishable from that of the wild type by epifluorescence imaging, with discrete membrane-associated patterns (Fig. Sup 2A), while western blotting showed wild-type levels of both mutated proteins (Fig. Sup. 2C). Both mutant *mreB* genes were cloned into an expression host. The corresponding proteins, purified to homogeneity (Fig. Sup 3A), displayed wild-type CD profiles (Fig. 3A). Unlike the corresponding *C. crescentus* mutant, both purified variants retained full polymerization proficiency *in vitro* under our sqTEM conditions, and no isolated single protofilaments were detected (Fig. 3B; Fig. Sup 3E). Both proteins also retained wild-type-like ATP affinity (Fig. 3E; Fig. Sup 3D), while V^114^A displayed near-wild-type ATPase activity (Fig. 3C; Fig. Sup 3B). By contrast, V^114^E displayed a significantly increased ATP hydrolysis rate (Fig. 3C). *B. subtilis* cells carrying this mutation displays only very mild shape defects (Fig. Sup 2A, B). We therefore asked whether the increased ATPase activity was associated with more subtle changes in MreB dynamics. We took advantage of the apparently preserved localization of these mutants to analyze their dynamics by total internal reflection fluorescence (TIRF) microscopy. This technique allows to observe proteins associated or embedded in the membrane and has previously provided insights into the dynamics of MreB foci, including their density and speed ^8, 9, 10^. The total density of MreB foci (Fig. Sup. 2F), the density of foci displaying directed motion (Fig. sup. 2E) –presumed to reflect the subpopulation of polymers actively involved in CW synthesis– and their speed (Fig. Sup. 2D) appeared unaffected by mutation V^114^A. For comparison, a strain carrying mutation E^136^A also displayed only marginal changes in MreB dynamics and foci density under our conditions (Fig. Sup 2D-F). In mutant V^114^E, however, the directionally moving foci displayed a modest increase in speed together with decreased density (Fig. Sup 2D, E). The mutation V^114^E may therefore partially destabilize MreB polymers or their association with Rod complexes, thereby reducing the fraction of directionally moving ‘active’ foci. However, whether and how the altered ATP hydrolysis, foci density and speed are mechanistically coupled remains unclear.

We next turned to the final selected mutant, G^56^R. This mutation introduced a large side chain predicted to clash with the following monomer along the protofilament (Fig. 2B; Sup. Mat.). G^56^R was therefore expected to significantly impair polymerization. Indeed, no polymer were detected by sqTEM (Fig. 3B), consistent with the *in vivo* data (Fig. 2). The purified protein retained wild-type ATP affinity (Fig. 3E; Sup 3D), indicating a preserved catalytic pocket, consistent with its wild-type CD profile (Fig. 3A). Unexpectedly, its specific ATPase activity was virtually extinguished with activity levels similar to that of the catalytic cleft mutants G^14^E and G^160^R (Fig. 3C; Sup. Table 4). This activity was also significantly lower than that of wild-type MreB in the absence of lipids (Fig. 3C), under conditions in which no polymers are detected by sqTEM ^12^. More specifically, under these non-polymerizing conditions, wild-type MreB retained approximately 50% of the specific ATPase activity measured under polymerization conditions in the presence of lipids (Fig. 3C). We previously described this unexplained activity as a futile ATP-hydrolysis cycle ^12^. Because G^56^R lies at the longitudinal monomer interface, ∼18 Å from the ATP γ-phosphate, it is unlikely to directly impair ATP hydrolysis or subsequent P_i_ release from the catalytic pocket (Fig. 2B). Its strongly reduced ATPase activity, measured by quantifying P_i_ release, therefore indicates that efficient ATP hydrolysis requires productive longitudinal MreB–MreB interactions, *i.e.*, contacts properly organized to support the conformational changes and reversible assembly dynamics of MreB, as well as the associated catalysis. The distinct effects of V^114^A/E on ATPase activity further suggest that perturbations primarily affecting lateral interactions may propagate through the assembly and modulate catalysis at the nucleotide-binding site, supporting the importance of an appropriate overall arrangement and/or dynamics of MreB–MreB contacts for efficient ATP hydrolysis. Moreover, these findings suggest that the apparent futile ATPase activity of wild-type MreB under non-polymerizing conditions may arise from transient intermolecular MreB-MreB contacts.

### High level adsorption to lipids depends on polymer formation

We next wondered how these mutations affect MreB/lipids interactions. We previously observed that wild-type MreB association with lipids was severely reduced in the presence of a non-hydrolysable ATP analog compared with ATP, leading us to conclude that ATP hydrolysis was required for MreB/lipid interaction ^12^. However, E^136^A retained full polymerization capacity in the presence of lipids (Fig. 3B) despite its severely reduced ATPase activity (Fig. 3C), prompting us to revisit this conclusion. We recently showed that without nucleotide, wild-type MreB has a limited intrinsic affinity for lipids, when it remains mainly monomeric, whereas lipid affinity increased strongly in the presence of ATP under polymerization conditions ^32^. To investigate MreB/lipid interactions, we turned to quartz crystal microbalance with dissipation measurements (QCM-D) to more accurately quantify the adsorption of wild-type and mutant MreB to supported lipid bilayers (SLBs) composed of an 80:20 mixture of DOPC:DOPG (see M&M section; Fig. Sup 4). QCM-D measures changes in resonance frequency and dissipation of a quartz crystal surface caused by interactions with an introduced compound. In these experiments, we recorded the absolute variation of frequency (∣Δf∣) of a SLB-covered crystal caused by the presence of MreB proteins, wild type or mutants. Here, the ∣Δf∣ increased after MreB injection (Fig. Sup. 4, arrow 6), reflecting the protein adsorption to the lipids. The maximal ∣Δf∣ reached after injection is hereafter termed Δf_adsorption_. Upon rinsing (Fig. Sup. 4, arrow 7), ∣Δf∣ decreased, consistent with the removal of adsorbed material from the SLB. The ∣Δf∣ remaining at the post-wash plateau, relative to the pre-injection baseline, is termed Δf_retention_ and indicates that some MreB material remained associated with the SLB under the rinsing conditions.

In a control experiment without nucleotide, wild-type MreB produced only a low |Δf| (< 10 Hz) that remained constant after washing and reflected limited basal adsorption of MreB to the SLB (Fig. 4A). Compared with the nucleotide-free condition, ATP-bound wild type MreB showed a large and rapid adsorption, as well as a rapid reduction following washing (Fig. 4A), consistent with enhanced association with the SLB. With ADP, adsorption was reduced, indicating weaker association with the SLB. Next, we examined the adsorption behavior of MreB mutants (Fig. 4B). Mutants G^14^E, G^56^R and G^160^R displayed a Δ*f*_adsorption_/Δ*f*_retention_ ratio close to 1, associated with low |Δ*f*| values (< 10 Hz) and little difference between ATP and ADP conditions, suggesting limited nucleotide-dependent adsorption to the SLB (Fig 4B). By contrast, the V^114^A, V^114^E, and E^136^A mutants displayed |Δ*f*| signals of 10-30 Hz in the presence of ATP with lower post-wash retention, and markedly reduced adsorption under ADP conditions, consistent with greater ATP-dependent adsorption (Fig 4B). Despite retaining polymerization proficiency, V^114^A and V^114^E exhibited altered ATP-dependent QCM-D responses (Fig. 4B): V^114^A adsorbed more slowly, whereas V^114^E showed lower adsorption and post-wash retention |Δf| values and a reduced Δ*f*_adsorption_/Δ*f*_retention_ ratio compared to wild type. Thus, perturbations predicted to affect lateral contacts also alter MreB association with the SLB (Fig. 4B). Altogether (Sup. Table 4), these results show that efficient MreB adsorption to SLBs does not require efficient ATP hydrolysis, because E^136^A polymerizes and adsorbs despite its severely (∼55-fold) reduced ATPase activity. More importantly, the E^136^A and G^56^R mutants reveal that ATP binding alone is insufficient to promote efficient MreB–lipid association. The proper overall arrangement and/or dynamics of MreB–MreB contacts (*i.e.* productive MreB–MreB contacts altered by G^56^R, V^114^A and V^114^E) and MreB ATPase activity (altered by E^136^A) appear to fine-tune the overall dynamics of polymer association with SLBs. Together, sqTEM and QCM-D data show that efficient ATP-dependent association with SLBs requires a polymerization-competent state.

**Figure 4.**
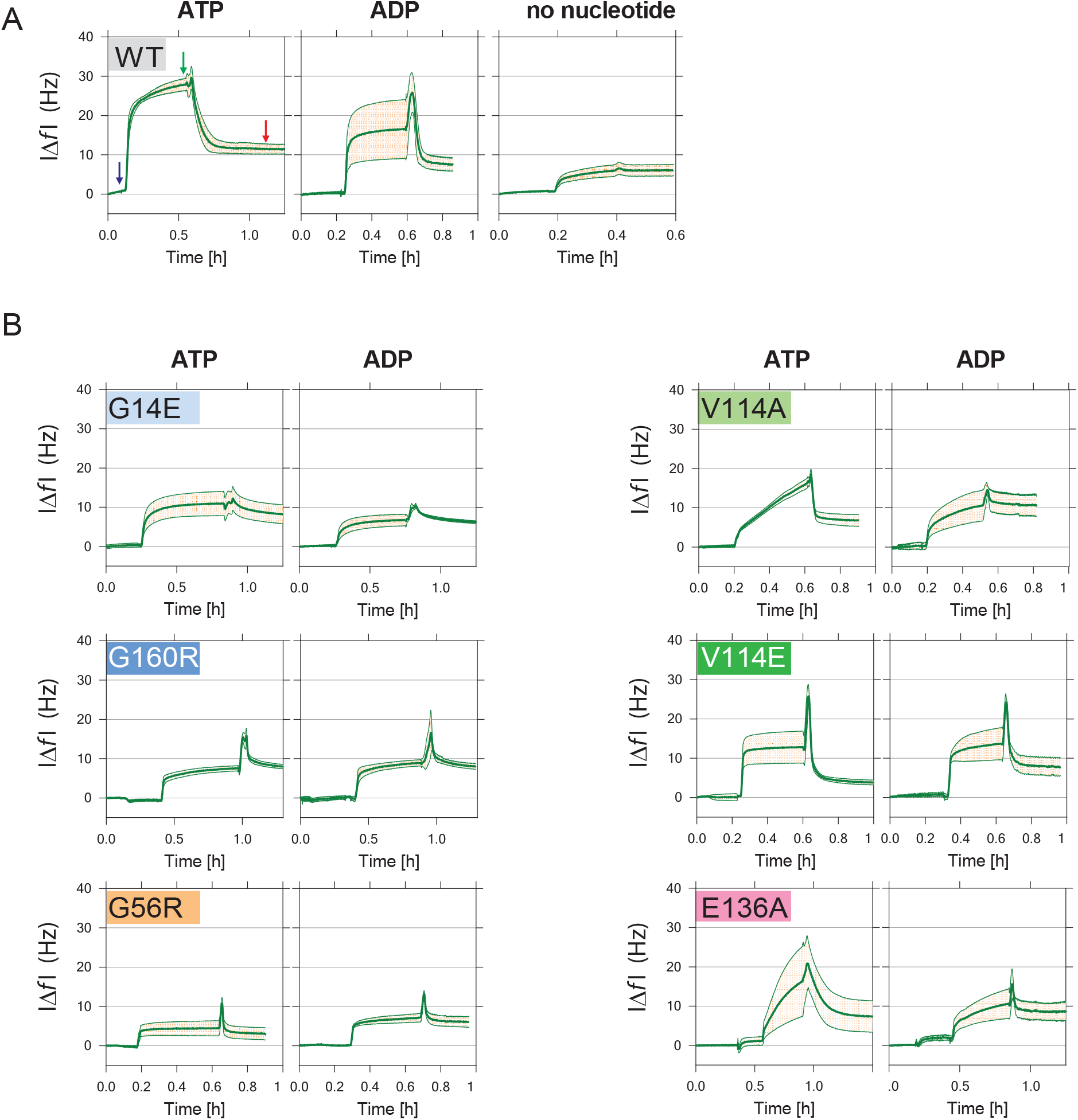
Ability to assemble into polymer predicts adsorption of MreB to a supported lipid bilayer. **A.** Kinetics of frequency changes (|Δ*f*|) in QCM-D experiments induced by addition of 1.3 µM of wild type MreB on an 80:20 DOPC:DOPG SLBs in the presence of 2 mM ATP or ADP or no nucleotide, and 2 mM of Mg^2+^. Arrows indicate : MreB injection (blue), wash (green) and post-wash |Δ*f*| plateau (retention). Standard deviation to the mean are displayed as colored areas. **B.** Kinetics of frequency changes (|Δ*f*|) in QCM-D experiments induced by addition of 1.3 µM of mutant MreB on an 80:20 DOPC:DOPG SLBs in the presence of 2 mM ATP or ADP, and 2 mM of Mg^2+^. Plots are averages of at least 2 independent experiments of 4 runs. Standard deviation to the mean are displayed as colored areas. Values are averages of at least 2 independent experiments of 4 runs. Error bars and standard deviation to the mean.

### ATP hydrolysis promotes disassembly of MreB polymers

Since a mutant with severely reduced catalytic activity (E^136^A) remains fully proficient in polymerization, we wondered about the role of ATP hydrolysis. It was tempting to draw a parallel with the actin model, in which P_i_ release following ATP hydrolysis promotes actin filament (F-actin) depolymerization ^36^. We therefore asked whether MreB might share this property to some extent with its eukaryotic homologue. We previously showed that, in the presence of limiting amounts of ATP, sqTEM reveals a two-phase process: an initial rapid formation of dense polymer fields (similar to our standard conditions with an excess of ATP), followed by a second, slower phase during which polymers disappear over 2-3 hours ^12^. Here, we used a similar setup to compare the disappearance of wild type and E^136^A MreB polymers. In a control experiment with excess ATP, 100 % of the fields were filled with a dense layer of polymers after 3 h for both wild-type MreB and the mutant E^136^A. When ATP was supplied in a limiting amount, the number of fields containing polymers of wild type MreB decreased markedly after 3h (Fig. 5A, B), as reported previously ^12^. By contrast, when the same experiment was performed with E^136^A, polymer disappearance was greatly reduced, with numerous fields remaining packed with polymers after 3 h. This suggests that ATP hydrolysis, as in F-actin, promotes MreB filament depolymerization. During our TEM acquisitions, we also noticed that while polymers disappeared over time under limiting ATP conditions, ghost-like structures remained visible at lower magnification (Fig. Sup. 5). This observation suggests that clusters of MreB may remain locally anchored to the lipids after the filaments have lost their ordered polymeric organization. Polymer disassembly may therefore be at least partly uncoupled from the resolubilization of MreB.

**Figure 5.**
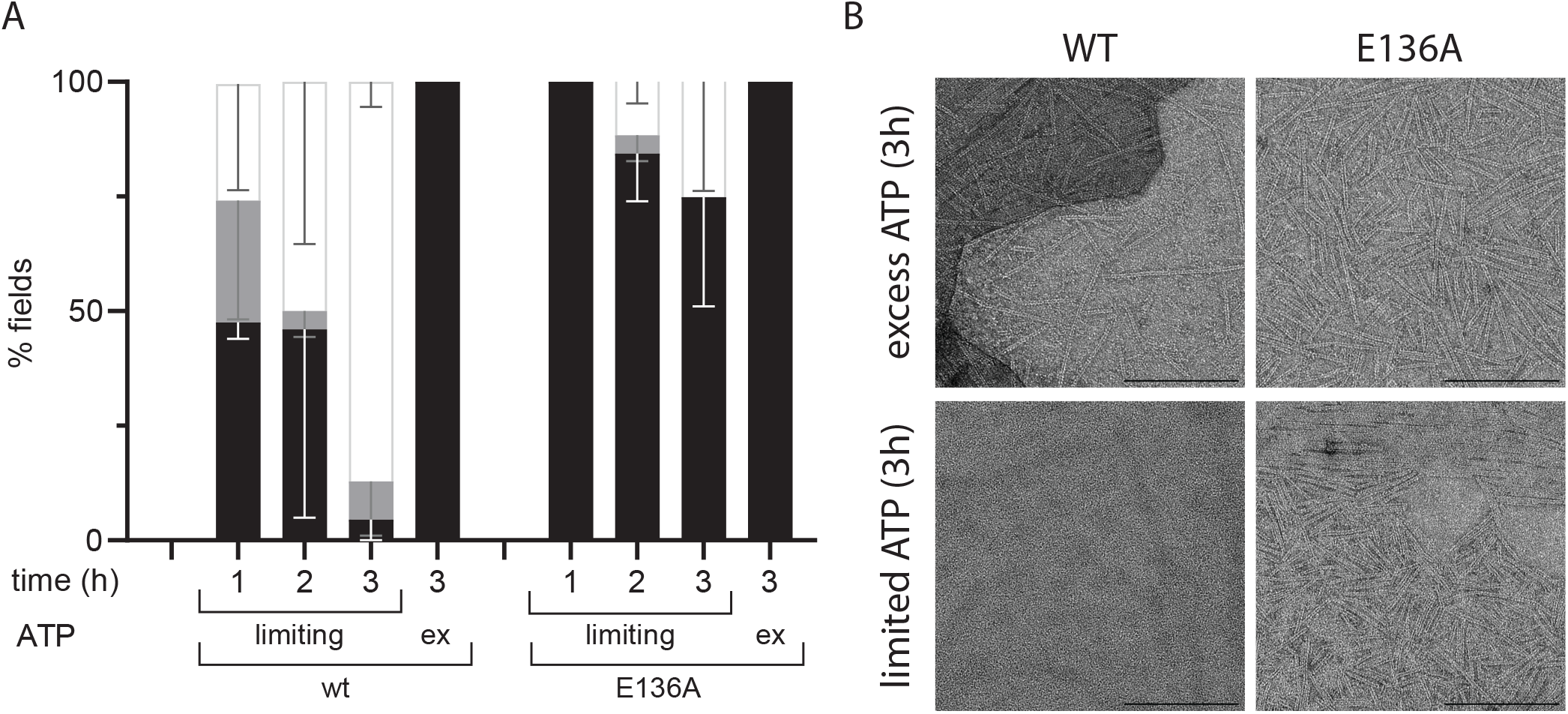
ATP hydrolysis promotes disassembly of MreB polymers. **A.** Mutation E^136^A stabilizes MreB polymers. sqTEM analysis, showing the frequency and density of polymer formation observed on negative stained EM images, of purified wild type (left) and E^136^A mutant (right) of MreB. 1.3 µM of proteins were set to polymerize in standard conditions, *e.g*. in the presence of excess ATP (2 mM), or in limiting amount of ATP 13 µM. Samples were fixed for TEM observation after 1 to 3h incubation at 37°C. **B**. Typical TEM images of wild type (left panels) and mutant E^136^A (right panels) taken after 3h incubation in the presence of excess (upper panels) or limiting (lower panels) amount of ATP. Scale bars : 200 nm.

## Discussion

In this study, we: (i) identified a highly sensitive reporter of MreB activity, (ii) used it in a genetic screen to identify MreB-deficient mutants and residues critical for MreB function *in vivo* in the G(+) model bacterium *Bacillus subtilis,* and (iii) characterized the corresponding variants of the homologous *Geobacillus stearothermophilus* MreB *in vitro.* These complementary cellular, biochemical, and biophysical analyses reveal key mechanistic steps coordinating ATP-driven MreB dynamics and support a model for its assembly-disassembly cycle.

### *mreBH*, a highly sensitive reporter of CW homeostasis

To design our screen, we exploited our previous observation that MreBH accumulates in the absence of MreB ^28^. The *mreBH*-based transcriptional fusion proved to be a powerful reporter of MreB function, although the underlying mechanism remains unclear. *mreBH* expression is under the dual positive control of SigI, an alternative sigma factor, and the WalKR two-component system ^37, 38^. WalKR monitors the activity of D,L-endopeptidases essential for PG expansion through their cleavage products and adjusts endopeptidase expression accordingly ^39^. By contrast, SigI is gradually activated in response to reduced PG synthesis, possibly through detection of defects in the PG meshwork ^40^. Thus, these regulators monitor two key aspects of PG homeostasis: synthesis and degradation. *mreBH* induction in an *mreB* knockout therefore suggests that both processes are perturbed, consistent with MreB’s proposed role in organizing PG synthesis and its genetic link to LytE, one of the two main essential D,L-endopeptidases ^41^. Remarkably, reporter induction reflected the severity of morphological and MreB-dynamic defects, detecting even mutations with barely observable phenotypes. Notably, the wild-type *gfp-mreB* strain, although phenotypically indistinguishable from its parental strain, also displayed slight induction over time (Fig. Sup 2G). This sensitivity, presumably arising from the dual regulation by SigI and WalKR, makes P*_mreBH_*a potentially valuable reporter of both MreB activity and PG homeostasis.

Several genetic screens for MreB mutants have been conducted previously, mostly in G(-) models, selecting gain-of-function suppressors resistant to the MreB inhibitors A22, YodL or YisK, or fast-growing suppressors of RodZ loss ^20, 21, 22, 23, 24^. These studies identified MreB contact sites for these inhibitors but provided limited insight into the overall mechanism of its polymerization. A recent alanine-scanning study identified 27 residues important for morphogenetic control in *E. coli* ^26^. Only three were also identified in our screen, two with mild effects beyond reporter induction. A strength of our approach is its use of a sensitive functional selection, which identified both critical residues and disruptive substitutions beyond alanine. However, many initially selected mutations could not be reconstructed, suggesting incompatibility with *B. subtilis* viability. Nevertheless, several substitutions caused a pronounced loss of MreB function (Sup. Table 4).

### On the respective roles of binding, hydrolysis of nucleotides and their coupling to polymer dynamics

Although catalytic-site mutations were expected from the loss-of-function screen, our integrated *in vivo* and *in vitro* analyses clarify the distinct contributions of ATP and ADP binding to MreB dynamics and of ATP hydrolysis to MreB polymerization and turnover. Previous ITC measurements using a membrane-binding-deficient *C. crescentus* MreB construct showed micromolar affinities for both nucleotides, with an approximately fourfold preference for ATP ^13^. By contrast, full-length *G. stearothermophilus* MreB bound both nucleotides with tens-of-nanomolar nanomolar affinity, showed an approximately twofold preference for ADP, and displayed markedly distinct thermodynamic signatures for the two nucleotides (Fig. Sup. 3C). ATP binding was almost entirely enthalpy-driven, consistent with the more extensive and stable nucleotide-Mg²⁺-protein-water interaction network involving the γ-phosphate observed in MreB structures ^12, 13, 14^. By contrast, ADP binding had a larger favorable entropic contribution, potentially reflecting greater water or ion release and/or conformational freedom, which may favor filament turnover.

Mutations affecting the intra- and interprotofilament interfaces were also expected from a loss-of-function screen, although their biochemical and mechanistic consequences were not. The G^56^R substitution, predicted to disrupt longitudinal contacts between consecutive monomers and located more than 20 Å from the ATP γ-phosphate, strongly reduced ATP hydrolysis, as monitored by P_i_ release. A similar phenotype was reported for MreB5, one of several paralogs in the wall-less bacterium *Spiroplasma citri* ^14^. In that study, the K^57^A substitution (R^57^ in our G(+) models), designed to weaken longitudinal intraprotofilament interactions, partially impaired MreB5 polymerization while also reducing ATPase activity. This biochemical similarity supports the idea that productive longitudinal MreB–MreB contacts contribute to efficient ATP hydrolysis. Together, these observations suggest that long-range coupling between longitudinal assembly and ATP hydrolysis may be conserved among at least some evolutionarily distant MreB orthologs.

We also investigated the V^114^E substitution, which was predicted to perturb the largest lateral protofilament interface (Sup. Mat.) ^13^. Indeed, V^114^ contributes to a prominent symmetric cluster of hydrophobic contacts within the extended network of MreB–MreB interactions between antiparallel protofilaments (Sup. Fig. 2H and Sup. Mat.). Unexpectedly, V^114^E had little effect on MreB polymerization *in vitro* but increased ATP hydrolysis and altered MreB-dependent dynamics *in vivo*. Whereas the equivalent substitution V^118^E in *C. crescentus* MreB disrupts lateral protofilament pairing *in vitro*, the *G. stearothermophilus* V^114^E variant formed paired protofilaments as efficiently as wild-type MreB, with no detectable single protofilaments under our conditions. Sequence differences within the *G. stearothermophilus* lateral protofilament interface may better accommodate a negatively charged residue at this position. For example, *G. stearothermophilu*s A^80^ replaces E^86^ in *C. crescentus* and may reduce local steric constraints. The increased ATP-hydrolysis rate of V^114^E was unexpected because its polymerization properties appeared wild-type-like. As V^114^ lies approximately 18 Å from the ATP γ-phosphate, perturbations of this lateral protofilament interface may allosterically modulate ATP hydrolysis, although the underlying mechanism remains unknown. One possibility is that V^114^E subtly destabilizes lateral protofilament interactions, accelerating polymer assembly–disassembly cycling and thereby increasing population-wide ATP turnover without measurably altering the overall polymer pool under our *in vitro* conditions. Intriguingly, V^114^E accelerated MreB-foci movements *in vivo*. Because MreB motion is driven by processive PG synthesis by the Rod complex rather than by MreB ATP-driven self-assembly, increased ATPase activity is unlikely to directly account for this phenotype. However, reduced lateral protofilament stability might explain the decreased number of active MreB foci. Their increased velocity might compensate for their reduced number, thereby helping to maintain the overall PG-synthesis rate, although this model remains speculative.

### A model for MreB assembly and disassembly cycle

Taken together, our study defines key requirements spanning the complete MreB assembly–disassembly cycle (Fig. 6, Sup Table 4). First, ATP binding, but not hydrolysis, generates a polymerization-competent MreB conformation, as shown by mutants G^14^E and G^160^R (defective for ATP-binding and polymerization) and E^136^A (defectives for ATP catalysis but retaining wild-type ATP binding and polymerization) (Fig. 3B,C,E). Second, productive longitudinal MreB–MreB interactions are required for efficient ATP hydrolysis, as illustrated by G^56^R. The mutant, impaired for longitudinal contacts, retains wild-type-like ATP binding but strongly reduced ATP hydrolysis and polymerization (Fig. 3B-C). Because wild-type MreB still hydrolyses ATP in the absence of lipids, thus under non-polymerizing conditions (Fig. 3C), this activity implies transient MreB–MreB interactions. It is tempting to postulate that small or short-lived oligomers resembling the nucleation step described for actin could form at this stage. Third, efficient association to lipids requires polymerization-competent MreB, rather than ATP binding or hydrolysis alone: G^56^R retains only basal lipid association despite wild-type ATP binding, whereas E^136^A polymerizes despite strongly impaired catalysis (Figs. 3 & 4). The contrasting lipid association profiles of polymerization-competent (E^136^A, V^114^A, and V^114^E) and -defective (G^14^E, G^160^R, and G^56^R; Fig. 4B) variants support a model in which polymerization enhances membrane association through assembly-dependent avidity, whereby multiple weak membrane contacts become jointly stabilized within MreB polymers, as also suggested for another morphogenetic protein, the bactofilin BacA^42^. Finally, ATP hydrolysis promotes filament destabilization, as demonstrated by the prolonged persistence of E^136^A polymers under ATP-limiting conditions (Fig. 5). Together, these findings support a comprehensive model of MreB assembly-disassembly cycle (Fig 6). ATP binding to monomeric MreB initiates the cycle. P_i_ release detected for wild-type MreB without lipids (Fig.6, dashed pathway) may represent a futile cycle driven by transient, unproductive MreB-MreB interactions in the absence of membrane. Membrane association may stabilize these contacts, while nucleation and assembly may reciprocally enhance membrane affinity by aligning and combining multiple weak membrane contacts within the growing polymer. This would promote elongation of the strongly membrane-associated polymers. ATP hydrolysis along the filament would then promote their destabilization and subsequent disassembly.

**Figure 6.**
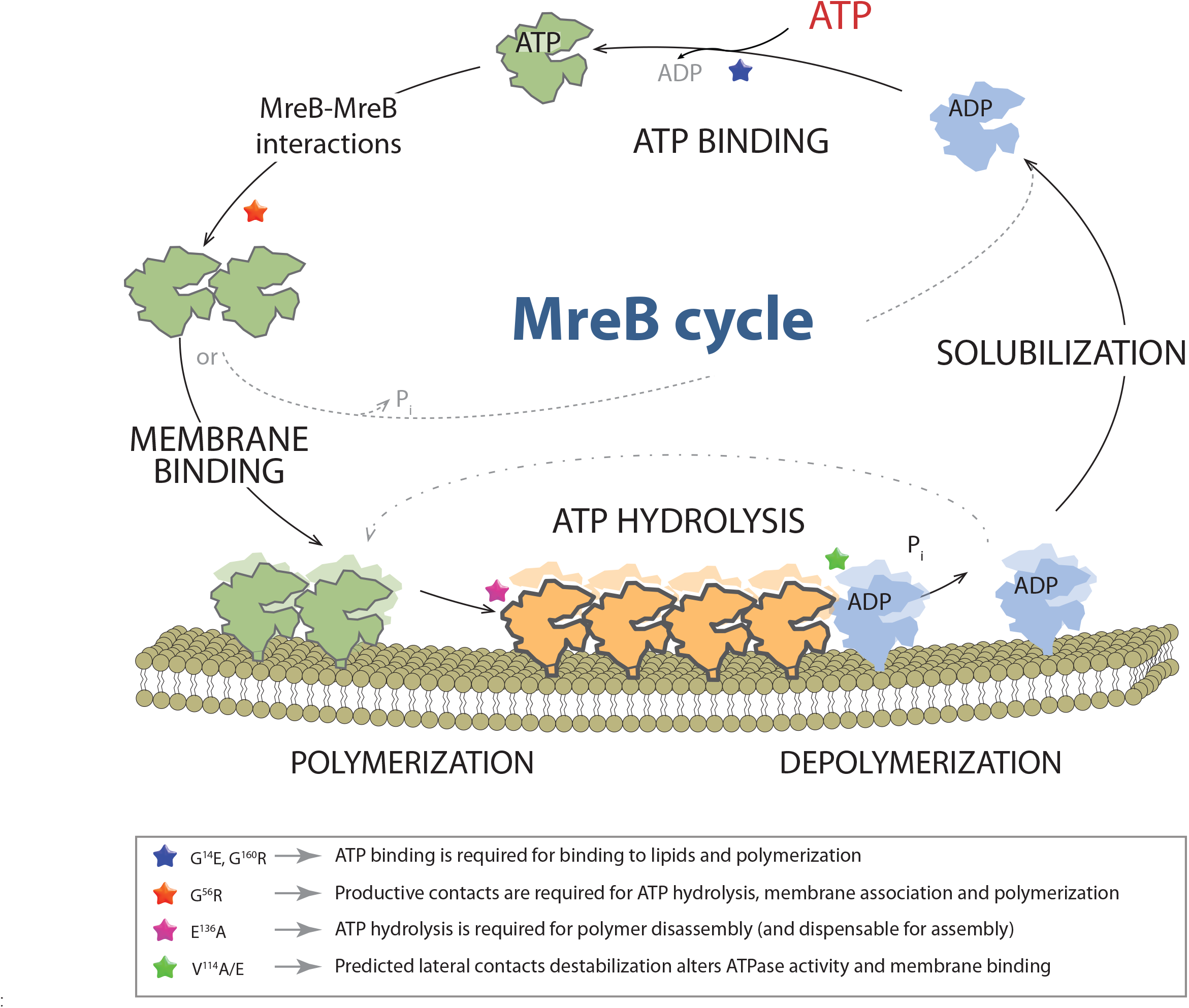
Model for ATP-driven MreB assembly–disassembly cycle. ATP binding generates a polymerization-competent MreB state and enables productive longitudinal contacts; ATP hydrolysis is not required for polymerization. Transient oligomerization/nucleation is proposed to precede membrane-bound polymer formation. Membrane association may stabilize MreB–MreB contacts, while assembly may reciprocally enhance membrane avidity by aligning multiple weak lipid-binding elements. ATP is hydrolyzed within membrane-associated polymers, promoting their destabilization and disassembly. ADP-bound subunits may remain transiently membrane-associated after loss of polymer order before their lower lipid affinity favors release into the cytosol. If ADP-to-ATP exchange occurs before release, membrane-associated MreB may re-enter assembly without complete solubilization (dash-dotted arrow). In the absence of lipids (broken line), MreB-MreB interactions induce low level ATP hydrolysis (futile cycle). Colored stars position the analyzed variants at the steps that they functionally probe. ATP-binding-defective G^14^E and G^160^R show limited binding to lipids, fail to polymerize and cause severe loss-of-function phenotypes *in vivo* (blue). G^56^R retains ATP binding but displays limited lipid interaction and disrupted polymerization, strongly reduced ATP hydrolysis, and impaired MreB function *in vivo*, supporting the coupling of productive longitudinal contacts with catalysis and membrane association (orange). The futile ATPase activity in absence of lipids (broken line) is strongly reduced by G^56^R, consistent with transient longitudinal MreB–MreB contacts being required for efficient hydrolysis. The hydrolysis-defective E^136^A variant retains strong ATP binding and membrane association, wild-type polymerization but delayed disassembly (magenta). V^114^A/E, predicted to be weakened for lateral interprotofilament contacts, remain polymerization-proficient but show altered membrane association; mutation V^114^E also increases ATPase activity and produces fewer, faster directionally moving MreB foci *in vivo* (green).

Under limiting ATP, ‘ghost-like’ structures observed by TEM (Fig. Sup. 5) suggest that polymers first lose cohesion while the proteins remain associated with the bilayer. The intermediate lipid association of ADP-bound MreB (between those of ATP-bound and nucleotide-free MreB; Fig. 4) is consistent with its temporary membrane retention after hydrolysis and may explain the slow disorganization of the polymers. This supports a model in which the re-solubilization of the protein would be delayed after MreB depolymerization. In this framework, if nucleotide exchange occurs before MreB release from the membrane, monomers may re-enter the cycle without fully dissociating from the membrane (Fig. 6; dash-dotted pathway), provided that the cellular metabolic state allows it. This prolonged lipid association may maintain a high local MreB concentration and favor polymer formation. Complementarily, in a related study, we observed nucleotide exchange within lipid-bound MreB filaments, indicating that ADP-to-ATP exchange can occur without prior depolymerization ^17^. MreB filaments may therefore be long-lived structures whose stability and remodeling respond to cellular nucleotide availability, although the energetic cost of this mechanism remains unknown.

In conclusion, our results support an MreB polymerization cycle that shares broad features with actin, including nucleation, polymerization, and filament destabilization following ATP hydrolysis. However, the two systems also differ substantially, notably in the strong coupling of MreB assembly to membranes, the lack of intrinsic MreB filament polarity ^17^, the central role of accessory proteins in regulating actin dynamics, and their distinct ATP/ADP binding properties, likely reflecting fundamentally different strategies. Whereas filament polarity and numerous actin-binding proteins coordinate actin dynamics by controlling the monomer pool and higher-order polymer organization ^36^, MreB polymers may remain stably anchored to the cell membrane for extended periods while energy remains available. This may reflect MreB’s direct coupling to and primary function as an anchor or organizing platform for the Rod complex PG synthesis machinery.

A key future challenge is to determine how nucleotide state and hydrolysis, membrane association, MreB conformational changes, filament architecture and dynamics, and Rod-complex interactions are integrated across molecular, structural, and cellular scales.

Resolving this integration will further clarify how ATP-driven MreB self-assembly provides a dynamic organizing platform for coordinating processive PG synthesis while remaining highly responsive to changes in cellular metabolic conditions.

## Materials and methods

### General methods and bacterial growth conditions

Methods for growth of *B. subtilis*, transformation, selection of transformants and so on have been described extensively elsewhere ^43^. DNA manipulations were carried out by standard methods, and amplifications performed with a high-fidelity DNA polymerase (Phusion, NEB), except for random mutagenesis (Taq, Takara Bio). All DNA amplicons were sequenced (Eurofins Genomics) following either their subcloning, for plasmids, or their direct integration into the *B. subtilis* host, for direct transformation of PCR, OE-PCR and isothermal assemblies (Gibson).

Isothermal assemblies were performed mainly as described in Gibson *et al*. 2009 with minor modifications ^44^. In brief, PCRs were generated with high fidelity Phusion (New-England Biolabs, NE, USA) DNA polymerase, and treated with *Dpn*I restriction enzyme to eliminate methylated DNA template. Equivalent quantities of purified PCR-generated DNA molecules were incubated with a joining mix (5 % w/v PEG-8000, 100 mM Tris-HCl pH7.5, 10 mM MgCl_2_, 10 mM DTT, 1 mM NAD, 0.2 mM each dNTP, 25 U/mL T5 exonuclease, 50 U/mL Phusion DNA polymerase, 6.7 U/µL Taq DNA ligase) and incubated 20 min at 50°C to form a single DNA molecule. 1 µL of the joining reaction was used as a template for PCR with external primers, and the resulting product was used to transform competent *B. subtilis* cells.

Quickchange mutagenesis on plasmids was performed by amplifying a template vector with two overlapping oligonucleotides introducing the change (mutation, deletion or insertion), followed by a dialysis of the product and digestion with the *Dpn*I restriction enzyme to remove template DNA, before transformation into *E. coli* for sub-cloning or into *B. subtilis*.

*Escherichia coli* strains were grown in LB medium and transformed with standard procedures with ampicillin (100 µg.ml^-1^) or kanamycin (25 µg.ml^-1^) selection. All *B. subtilis* are derivatives of the laboratory collection strain 168 and were grown 37 °C in rich lysogeny broth (LB) medium supplemented with 20 mM MgSO_4_ and/or antibiotics when indicated. Antibiotics were used at the following concentrations: chloramphenicol, 5 μg/ml; kanamycin, 10 μg/ml; spectinomycin, 100 μg/ml; erythromycin, 10 μg/ml; neomycin, 15 µg/ml. *B. subtilis* strains, plasmids and oligonucleotides used in this study are listed in Sup. Table 1, 2 and 3 respectively.

### Strains and plasmids constructions

#### Construction of a strain carrying a neomycin resistance cassette upstream of the *mreB* operon RCL414 (ABS2149)

A 2416 bp DNA fragment containing a fraction of *maf*, *radC* and the neomycin resistance cassette cloned upstream of the proximal promoter of the *mreBCD* operon, was PCR amplified using as a template the chromosomal DNA of strain 3725, otherwise deleted for *mreB*, and the oligonucleotides ac1345/ac1346. A second 1726 bp DNA fragment containing *mreB* and a fraction of *mreC* was produced using a derivative of the 168 strain and ac1334/ac1335 as primers. The purified products, overlapping by 44 bp, were next used as templates together with the external ac1345/ac1335 in an OE-PCR to generate a 4098 bp, and the final product was transformed into the recipient wild type 168 *B. subtilis* strain. Recombinant clones were selected on LB plates supplemented with kanamycin and 20 mM magnesium.

#### Construction of a strain expressing P*_mreBH_ lacZ* (RCL422)

A transcriptional fusion between the promoter region of *mreBH* and the *lacZ* reporter gene from *E. coli* was cloned in *B. subtilis* at the ectopic *thrC* locus. For this, a 299 bp DNA fragment was PCR-amplified from 168 chromosomal DNA using the oligonucleotides ac1246/ac1247, extending from the upstream *abh* gene to a putative ribosome-binding sequence (rbs) directly preceding the *mreBH* start codon. The fragment was digested and cloned into the corresponding *Eco*RI/*Bam*HI restriction sites of pDG1729, generating pAC783, and the resulting suicide vector transformed into *B. subtilis* 168, generating ABS1755. Integration events were selected with spectinomycin and checked for proper integration by PCR. A derivative strain was constructed by transforming ABS1755 with pDag32, allowing the disruption-replacement of *spc* with a chloramphenicol resistance cassette (*cat*).

#### Construction of a strain expressing P*_nat (o-r)_gfp^A206K^mreB* (RCL421)

We previously reported that *gfp-mreB* fusions expressed under an inducible promoter lead to over-accumulation of the protein during the transition phase ^8^, but that if the same fusion was placed under its natural promoter, the protein levels were lower than that of the wild type during exponential growth ^28^, presumably because of a lower translational efficiency of *gfp*. We constructed a minimally perturbative fusion overcoming both issues by placing at its natural locus (thus keeping all the natural transcriptional regulations) a *gfp-mreB* fusion, preceded by an optimized rbs (o.r) to enhance translation ^45^. We also used a monomerized variant of GFP (A206K) to minimize the risk of localization artefact ^46^. For this, we performed isothermal assembly of two PCR-generated DNA fragments. One was amplified with oligonucleotides cc181/cc186 using chromosomal DNA from RCL414 as a template and carrying the region upstream of *mreB* and a *neo* cassette, and one with cc182/cc185 as primers and DNA from RCL238 carrying a *gfp-mreB* translational fusion. Overlapping cc186 and cc185 introduced the optimized rbs we designed according to the work from Vellanoweth and coworkers. The assembled fragment was subsequently transformed into 168 *B. subtilis* strain, generating strain RCL420. We next introduced the mutation A206K in GFP by similarly transforming an isothermal assembly product of 2 PCR-generated DNA fragments: one amplified with oligonucleotides cc181/ac1282 and one with cc182/ac1281 using RCL420 as DNA template, the overlapping primers ac1281-ac1282 bearing the nucleotide change. The product was amplified by PCR using primers cc181/cc182 and transformed into wild type 168 *B. subtilis* generating the final strain RCL421 carrying P*_nat (o.r)_gfp^A206K^mreB*.

#### Construction of *E. coli* strains for expression and purification of recombinant MreB

All plasmids to produce mutated version of MreB from *G. stearothermophilus* are derivatives of pCC110 ^12^, and were generated by quickchange mutagenesis. The resulting plasmids (listed into Sup. Table 2) were transformed into the “T7 express” expression strain of *E. coli*.

### Random mutagenesis and selection for *mreB* mutants of interest

Low fidelity PCR was performed with Taq polymerase in the presence of 0.4 mM MnCl_2_ in order to enhance the mutation rate, using oligonucleotides ac1345/ac1335, and chromosomal DNA from strain RCL414 as template (Fig. Sup. 1B). RCL414 carries a neomycin resistance marker upstream of the *mreBCD* operon. The PCR product was dialyzed and transformed into naturally competent *B. subtilis* RCL422, carrying the P*_mreBH_lacZ* reporter. Cells were plated on LB-agar supplemented with neomycin for the selection of integration events, X-gal for screening of MreB deficient clones, and Mg^2+^ to ensure their survival (Formstone, 05). Blue clones were isolated and the area that was PCR-amplified prior to transformation (from 3’ of *maf* to 5’ of *mreC*) entirely sequenced. Clones carrying a frameshift mutation, a stop codon or no mutation in the *mreB* orf were discarded. The screen revealed 51 different amino-acid changes in MreB.

### Construction of strains carrying mutated *gfp-mreB* fusions by site-directed mutagenesis

We successfully constructed 33 strains carrying a mutant *mreB* gene at its natural locus in fusion with *gfp*, each bearing a unique SNP identified in our screen. For this, we used isothermal ‘Gibson’ assembly to combine PCR fragments generated on DNA template from strain RCL421 carrying a P*_nat (o.r)_gfp^A206K^mreB* fusion (Fig. Sup. 1D). The two PCR products were generated using cc181 or rk14 as external oligonucleotides and two (overlapping) forward and reverse internal primers specific of each mutation to introduce (see Table Sup. 3). Gibson-generated DNA fragments were either directly transformed or amplified by PCR using cc181/rk14 before transformation into competent *B. subtilis* cells of strain RCL422, and selected on LB plates supplemented with neomycin and Mg^2+^. Resistant clones were isolated and the presence of the *gfp-mreB* fragment verified by PCR. Finally, the complete region from *maf* to *mreC* was sequenced to confirm the presence of the mutation and the absence of putative additional SNPs, as a consequence of the amplification/assembling process.

### Epifluorescence and TIRF microscopy acquisitions

Images and movies were taken on at least two different days for each strain. Sampling were done on exponentially growing cell at 37°C in rich LB medium without antibiotic or Mg^2+^ supplement. For epifluorescence microscopy, samples were mixed with the membrane dye Nile Red (final concentration 10 µg/ml), before spotting on a 2 % agarose-LB pad as previously described ^47^. Phase contrast and epifluorescence (Em. filter 630 nm) images of the stained membranes were acquired on an inverted Nikon Ti-E microscope equipped with a 100x CFI Plan Fluor objective (NA 1.3, WD : 0.2 mm) and an iLas2 laser coupling system from Roper Scientific (50 mW, 561 nm) and an Orca-R² Hamamatsu CMOS camera with a final pixel size of 64.5 nm. Bright field and TIRF acquisition were acquired similarly except that no dye was used, the acquisitions were collected with an electron-multiplying charge-coupled device (EMCCD) camera (iXON3 DU-897, Andor) at maximum gain setting (300) attached to a 2.5x magnification lens, a 100x Apochromat (NA 1.49) objective, and an azimuthal TIRF under iLas2 laser coupling system from Roper Scientific (150 mW, 488 nm). Final pixel size was 64 nm. All images acquisitions were controlled by Metamorph v.7 software. Cell width determination were performed on epifluorescence images of membrane-labelled cells with the MicrobeJ plugin ^48^) on Fiji software ^49^, and calculated as the mean of the width along the median axis. For determination of particle dynamics: image segmentations, single particle detection and tracking (SPT) along the time lapses, and trajectory classification by mean squared displacement (MSD) analysis were performed as previously described ^50^.

### Western Blot

Strains were grown in LB medium supplemented with 20 mM MgSO_4_ at 37 °C until OD_600 nm_ = 0.4. 2 mL of cultures were spun down at 13000 rpm for 5 min at 4°C and frozen at −20 °C until further use. The pellets were resuspended in 25 µL resuspension buffer (50 mM glucose, 1 mM EDTA, 50 mM Tris pH 8, 10 mg/mL lysozyme, cOmplete™ protease inhibitor cocktail (Roche, Merck)) and incubated for 5 min at room temperature. Then, 25 µL of ice-cold lysis buffer (500 mM NaCl, 1 % NP40, 50 mM tris pH 8, 5 mM MgCl_2_, 0,05 % benzonase®) were added, mixed and incubated on ice for 15 min. 12 µL of samples were mixed with 3 µL of 5x loading dye (250 mM Tris pH 6,8, 8% SDS, 40% Glycerol, 0.02 % Bromophenol blue, 0,04% β-mercaptoethanol), heated at 95°C for 5 min, and loaded on a Mini-PROTEAN® TGX™ precast gel (BIO RAD) and migrated for 45 minutes at 150 V in 1x Tris-Glycine-SDS Buffer (EUROMEDEX). Next, the proteins were transferred with an iBlot™ 2 dry blotting system (Invitrogen) on a NC regular stack (Invitrogen) using a 3-step sequence of 20, 23 and 25 V for 1, 4 and 2 min, respectively. The quality of the transfer was controlled using a Ponceau red staining solution. After briefly rinsing with water, the membrane was blocked with 5% milk in TBS-T (10 mM Tris pH 8, 150 mM NaCl, 0,05 % Tween 20) for 20 min. The membrane was incubated first with anti-MreB primary antibodies at 1/10000 in TBS-T buffer overnight, then with anti-rat HRP-conjugated secondary antibodies at 1/10000 in TBS-T for 2 h. Each incubation was followed by extensive washes with TBS-T. Membranes were revealed with the Clarity™ Western ECL Substrate (BioRad), and imaged on a ChemiDoc™ MP (BioRad) imaging system.

### Protein purification

Expression of recombinant MreB proteins and subsequent purifications were performed as previously described (Mao et al, 20). In brief, 6his-tagged proteins were produced in the expression host by overnight induction and cell were disrupted by sonication. After clarification, proteins were purified following a two-step procedure, first via affinity chromatography on a Ni-nitrilotriacetic acid (NTA)-agarose resin followed by size-exclusion chromatography (HiLoad 16/60 Superdex® 200 pg). Protein preparations were quantified, concentrated by ultrafiltration spin columns up to a maximum concentration of 14 µM, aliquoted and flash frozen for storage at −70°C until further use.

### Circular Dichroism

The secondary structure of recombinant wild type and mutant forms of purified MreB were analyzed by circular dichroism (CD). Proteins were dialyzed against of 10 mM NaPO_4_ pH 7 buffer containing 200 mM NaSO_4_. Far-UV spectra were recorded on a Jasco J-810 spectropolarimeter (Japan Spectroscopic Co., Tokyo, Japan at 20 °C in a 0.5 mm path-length quartz cuvette after the sample compartment was purged with nitrogen. Each CD spectrum was averaged over four scans, collected at a scan rate of 50 nm/min, recorded from 280 to 200 nm, on samples with peptide bond concentration ≥ 1.6 mM. Baseline spectra obtained with buffer were subtracted for all spectra.

### TEM & sqTEM

Polymerization efficiency was determined on transmission electron microscopy (TEM) images, as described previously (Mao 23). For this, 50 µg/ml (1.3 µM) MreB was set to polymerize in reaction buffer (20 mM Tris pH7, 100 mM KCl, 5 mM Mg^2+^, 2 mM ATP) on a 20 µl droplet topped with a monolayer of lipids. The monolayer results from dropping a 0.4 µl of *E. coli* polar lipid extract (#100600, Aventi Research™, Merck) on the reaction droplet. After 1h incubation at 37°C, a carbon-coated EM grid (CF300-cu, EM sciences), was deposited on top of the droplet for 2’, fixed and stained with a 2% uranyl acetate for 2’, and air dried. TEM images were acquired on a charge-coupled device 553 camera (AMT) on a Hitachi HT 7700 electron microscope operated at 80 kV (Milexia – France). For semi-quantitative TEM (sqTEM), we used the protocol previously described ^12^. Briefly, 12 positions widespread on the entire EM grid were predetermined, observations were made at high magnification on each of these precise localizations (no looking around for polymers) and polymer densities were distributed into 3 categories: loan, low density, or no polymers. Results are averages of several independent polymerizations performed on different days.

### ATPase assay

Release of free inorganic phosphate (Pi) was measured as a proxy for ATP hydrolytic activity, using the malachite green detection technic, as described in ^12^. For this, several mixes using a range of [MreB] were setup in a reaction buffer (20 mM Tris pH7, 100 mM KCl, 5 mM Mg^2+^, 0.5 mM ATP) containing 0.5 mg/ml liposomes. Liposomes were made fresh by resuspending desiccated aliquot of *E. coli* polar lipid extract into H_2_0 at 10 µg/µl. The reactions were initiated by the addition of MreB and transfer to 53°C, and ended after a 1h incubation by addition of 1 reaction volume of malachite revelation buffer (0.2 % (w/v) ammonium molybdate, 0.7 M HCl, 0.03 % (w/v) malachite green, 0.05 % (v/v) Triton X-100). [Pi] was determined by measuring the absorbance at 650_nm_ on a 96-well plate spectrophotometer (Synergy 2, Biotek) and blanked against a mock reaction devoid of protein.

### TSA

To remove traces of EDTA, protein samples were first buffer exchanged against a 20 mM Tris pH7, 100 mM KCl buffer using filtration spin columns (Vivaspin, 10 MWCO), concentrated and quantified. 2.5 µM of protein were added to reaction mixes containing 20 mM Tris pH7, 100 mM KCl, 10x Sypro ™ Orange (Invitrogen), and when indicated 2 mM MgCl_2_, and/or 50 µM of nucleotide (ATP, ADP or GTP), in a final volume of 30 µl. All the reactions were distributed in duplica on a Microamp® fast 96-well reaction plate (AppliedBiosystems, ThermoFischer Sc.) and subjected to 1°C / min temperature ramp from 25° to 95°C on a StepOnePlus thermocycler (AppliedBiosystems, ThermoFischer Sc.). Fluorescences were recorded and processed with StepOne™ software (v 2.3) and the T_m_ determined as the average peak fluorescence of melting curve’s derivative of the replica. Reported values are average of at least 2 independent experiments.

### Isothermal titration microcalorimetry

Measurements were performed with a PEAQ ITC isothermal titration calorimeter (Malvern Panalytical, Malvern, UK), at 25°C. Nineteen injections of 2 µL of ligand solution (ATP or ADP) at 90 µM in a reaction buffer (20 mM Tris pH7, 100 mM KCl, 2 mM MgCl_2_) were performed every 180 s in the chamber containing 200µl of protein at 20 µM in identical buffer, with rapid 500 rpm mixing. Theoretical titration curves were fitted to experimental data with PEAQ ITC software supplied by Malvern. This software uses the relationship between the heat generated by each injection and ΔH (enthalpy change in kcal.mol^−1^), Ka (the association binding constant in M^−1^), n (the number of binding sites), total protein concentration and free and total ligand concentrations.

### QCM-D analysis of MreB interaction with a supported lipid bilayer membrane

#### Lipid preparation

In a previous study, we identified supported lipid bilayers of DOPC and DOPG 80%:20% (v/v) as the optimal lipid mixture to study the MreB/lipid interaction by QCM-D (Mao et al, 2023). In this study, we kept this lipidic composition but used solvent-assisted lipid bilayer formation (SALB) method ^51^ to prepare the SLB bilayers. For this purpose, appropriate volumes of lipid stock in chloroform (DOPC, and DOPG; Avanti Polar Lipids, Merck) were added to a glass vial in the desired molar ratio. For our measurements, we required 1.5 mL total volume which amounts to a total amount of lipid of 0.75mg. The chloroform was subsequently evaporated using a gentle stream of argon while slowly rotating the glass vial at a slightly tilted angle. After the chloroform was visibly evaporated, the glass vials were placed into a vacuum oven overnight at room temperature. The lipids were dissolved in isopropanol just before being used at a final concentration of 0.5 mg/mL. The lipid solution was gently vortexed for 10s to ensure homogenization.

#### Quartz crystal microbalance with dissipation monitoring

QCM-D measurements are based on frequency and dissipation changes on a quartz crystal surface ^52^. Measurements were performed on a QCM-D E4 (QSense AB, Biolin Scientific AB, Gothenburg, Sweden) as reported in previous lipid bilayer-protein interactions and adhesion studies ^12, 53^. Custom-made quartz crystals (QSense AB, Biolin Scientific AB, Gothenburg, Sweden) were prepared as described previously ^12^ up to their transfer into the measurement chambers, set to 25°C. Peristaltic pumps were set up to inject HEPES–NaCl buffer (10 mM HEPES pH 5.5, 100 mM NaCl) into the chamber at a flow rate of 100 µL/min until the baseline was stable. After baseline stabilization, isopropanol was pumped into the measurement chambers at a flow rate of 100 µL/min, and after frequency and dissipation were stabilized we continued to flow isopropanol into the chamber for at least another 10 min to replace HEPES–NaCl buffer. After isopropanol had completely replaced HEPES–NaCl buffer, we added the 0.5 mg/mL DOPC:DOPG lipid mixture in isopropanol to the measurement chamber at a flow rate of 100 µL/min for at least 10 min until the measurement signals stabilize. This step usually resulted in frequency losses of ∼4-7 Hz due to the adsorption of lipids on the silicon dioxide surface. In the final solvent replacement step, HEPES–NaCl buffer was pumped into the system at a flow rate of 100 µL/min for 20 min in order to enforce spontaneous SLB formation. This step ended in frequency (−26-30 Hz) and dissipation (0.1-1×106) changes that indicated successful SLB formation. After another 20 min, protein buffer (1 mM MgCl_2_, 100 mM KCl, 10 mM HEPES (pH7.5)) was added to the SLB. Once a stable baseline is achieved, 1.3 µM of MreB protein in protein buffer (± 2 mM ATP, ADP, and no ATP) were added to the SLB at a flow rate of 100 µL/min for approximately 3-5 min, then the pump was stopped. The adsorption of MreB was measured for at least 20 min before exchanging and rinsing with MreB buffer at 100 µL/min. The analysis software QTools (QSense AB, Biolin Scientific AB, Gothenburg, Sweden) was used to extract raw data (kinetics, frequency and dissipation changes) and the data was analyzed using Graphpad Prism (GraphPad Software, USA). Each measurement was repeated at least twice with 4 repeats per run.

## Acknowledgments

This work benefited from the MIMA2 facility (Université Paris-Saclay, INRAE, AgroParisTech, GABI, 78350, Jouy-en-Josas, France) for TEM observations, the PIM platform of I2BC for TSA and ITC approaches, supported by French Infrastructure for Integrated Structural Biology (FRISBI) ANR-10-INBS-05, and the Biophysics technical platform at the VIM research unit (INRAe, VIM-UMR0892, Jouy-en-Josas, France) for CD acquisitions. We thank Magali Aumont-Nicaise, Drs Davy Martin and Human Rezaei for their precious help and fruitful discussions. We also thank past and present members of the ProCeD lab for their expertise and discussions, with a special emphasis to Cyrille Billaudeau.

---

**Figure S1.**

**A.** P*_mreBH_ lacZ* transcriptional reporter is induced in a *mreB* KO (RCL423) but not in a wild type (RCL422) strain of *B. subtilis*. A strain carrying *gfp* in translational fusion with *mreB* at its natural locus (RCL424) does not express the reporter gene. Exponentially growing cultures were spotted on LB medium plate supplemented with 20 mM MgSO_4_ and X-gal, and grown over night at 37°C.

**B.** Schematic of the two-step generation of mutants by PCR-mutagenesis.

**C.** Summary of the selection step.

**D.** Schematic of the 3-step construction of clean strains, each carrying a *gfp-mreB* fusion with a single point mutation, at its natural locus with optimized expression.

**E.** Multiple sequence alignment of MreB from *G. stearothermophilus* (G.ste), *B. subtilis* (B.sub), *T. maritima* (T.mar), *E. coli* (E.col) and *C. crescentus* (C.cre), using Clustal Ω. Conservation between sequences is indicated below as full (*), strong (:) or weak (.). Secondary structure information (α-helices and β-sheets) of *G. stearothermophilus* are displayed above the sequence based on ^12^, with colors referring to the four subdomains of the protein: IA (blue), IB (yellow), IIA (red) and IIB (green). The sequences corresponding to each subdomain are boxed accordingly with matching colors. Black boxes frame the hydrophobic residues involved in protein/lipid interactions ^12, 18^. Residues in contact with ATP are highlighted in purple, based on *G. stearothermophilus* MreB structure (PDB 7ZPU; Mao *et al*., 2023). Residues involved in lateral contact between protofilaments (red), and longitudinal contacts on subdomains A (yellow) or B (orange) are highlighted based on the crystal structure of A. *crescentus* MreB (PDB 4CZJ; Van den Ent *et al.*, 2014). Mutated residues isolated in *B. subtilis* during the screen are highlighted in cyan, and the ones mutated on *G. stearothermophilus* MreB in green.

**Figure S2.**

**A.** Microscopic observations of *B. subtilis* mutants. Cells were grown in rich LB medium to mid-exponential phase of growth, colored with the membrane stain Nile Red and imaged by phase contrast (phase), or in green (GFP) or red (membrane) fluorescence. Top row: mutants from the screen displaying morphological defect identical to an *MreB* KO strain and a complete loss of GFP-MreB localization, plus the control strain expressing wild type *gfp*-*mreB* (’’WT’’, RCL424). Bottom left: mutants with a mild phenotype. Bottom center: mutant from the literature designed to alter the catalytic ATPase activity of MreB. Bottom right: control strains deleted for *mreB* (’’Δ*mreB*’’; RCL423) and the parental wild type reference, devoid of insertion or *gfp* fusion to *mreB* (’’wt168’’; RCL422). Images displayed are typical results of at least two independent replica. **B**. Comparison of *B. subtilis* cell width from images on A. Data are a compilation of at least two independent experiments. Statistical significance between wild type and mutant’s width was calculated using the Welch’s non-parametric unpaired t-test (**= 0.001<*P*<0.01; *= 0.01<*P*<0.05; ns = *P*>0.05). **C.** Western blot analysis. Strains grown in rich medium to mid exponential growth were harvested, normalized by optical density, prior to total cell extractions, load on SDS-PAGE, transfer and blotted with anti-MreB antibodies. Wt (44) correspond to the parental wild type 168 strain and display a single band corresponding to MreB. Mutants deleted for *mreB* (413, 423) show a faint band due to cross-reacting species with the paralogous Mbl. All other strains present a *gfp* fusion inserted in frame with *mreB* at its natural locus (responsible for the size shit), and were either wild type (424) or mutant for *mreB* (8 right -hand lanes). Display is a typical result of two independent replica. **D-F**. Dynamic properties of GFP-MreB foci of the wild type (Wt) and mutant strains V^114^A, V^114^E and E^136^A. MreB dynamics were determined from TIRF microscopy acquisition on cells grown to mid-exponential phase in rich LB medium. Tracked foci during time courses were classified by MSD analysis. Data are compilations of at least two independent experiments, and individual values of each plot, the average per cell. Statistical significance between wild type and mutant’s was calculated using non-parametric nested t-test (*= 0.01<*P*<0.05; ns = *P*>0.05). **D**. Velocity of GFP-MreB foci displaying directional motion. **E.** Density of foci per cell surface displaying directional motion, as determined by MSD analysis. **F.** Total density of foci regardless of their dynamic properties. . **G**. Comparison of P*_mreBH_ lacZ* reporter induction in the presence of wild type or mutated *gfp-mreB* fusions. Upper row: parental wild type strain without the reporter (168), and strains with the reporter and either deleted for *mreB* (Δ*mreB*; RCL423) or carrying wild type *gfp*-*mreB* (wt; RCL424); lower row: strains carrying the reporter and a mutated version of *gfp-mreB*. **H.** Localization of the studied mutations in an AlphaFold-predicted model of six ATP–Mg²⁺-bound *G. stearothermophilus* MreB subunits arranged as two antiparallel three-subunit protofilaments. The model was predicted with high confidence (ipTM = 0.82; pTM = 0.84) and provides an alternative to the single-protofilament-like ATP-bound crystal structure shown in Fig. 2B (PDB 7ZPU; ^12^). Relative to Fig. 2B, movement of subdomain IB closes the IB–IIB cleft. The predicted model closely resembles the antiparallel protofilament arrangement of AMPPNP–Mg²⁺-bound *C. crescentus* MreB (PDB 4CZJ; ^13^), for which the longitudinal and lateral MreB–MreB interfaces are analyzed in the Sup. Mat.). Indeed, the model aligns more closely with PDB 4CZJ than with 7ZPU, with RMSDs of 0.77 versus 1.48 Å over 310 residues. Left: front view highlighting the contacts at or near longitudinal intraprotofilament interfaces. The protofilament is oriented and colored as in Fig. 2B, with MreB subdomains IA (pale blue), IB (pale yellow), IIA (pale red), and IIB (pale green). Right: side view, rotated 90° relative to the left view, showing lateral inter-protofilament contacts between the antiparallel protofilaments and the potential contribution of V114.

**Figure S3.**

**A.** Coomassie staining of SDS-polyacrylamide gel comparing the preparation of purified MreB proteins, wild type and mutants. Each lane contains 2 µg of pure protein. **B.** ATPase activities of wild type and mutants of MreB, estimated as the concentration of inorganic phosphate (Pi). P_i_ was determined by the malachite green assay after 2h incubation at 37°C in polymerization conditions, in the presence of a range of concentration of MreB. Values are averages of four independent experiments and error bars are standard deviations to the mean. **C.** Isothermal titration microcalorimetry comparing the binding of ATP and ADP to wild type MreB. Top panels show the temperature variation upon injection of ligand (ATP or ADP at 200 µM) to the chamber containing wild type MreB at 20 µM. Bottom panels show the integrated heat. Solid line is the best fit on a single binding model. Right panels are the control experiment without protein showing the effect of the buffer and ligand mix injected to the chamber. All experiments were performed in the presence of 2 mM Mg^2+^. **D.** Compared melting temperature (Tm) of wild type (WT) and mutant of MreB, determined by TSA, in the presence or absence of ATP. Tm were recorded in the presence of 2.5 µM of protein, 50 µM ATP and 2 mM Mg^2+^. Values are averages of at least two independent experiments and error bars are standard deviations to the mean. 1. **E.** TEM images of wild type (WT) and mutant forms of MreB set to polymerize in the presence of ATP and lipids. All proteins form similar dual protofilaments. Scale cars: 200 nm.

**Figure Sup 4.**

Typical complete spectrum of a QCM-D experiments, recording the kinetics of frequency change (|Δ*f*|) over time. (1) HEPES–NaCl buffer (10 mM HEPES pH 5.5, 100 mM NaCl) is pumped into the measurement chamber until the baseline is stable. (2) Buffer is replaced by isopropanol, until 10’ after frequency and dissipation stabilize. (3) DOPC:DOPG lipid mixture (0.5 mg/mL in isopropanol) is added until 10’ after the measurement signals stabilize. (4) Solvent is replaced by HEPES–NaCl buffer enforcing SLB formation. (5) reaction buffer (1 mM MgCl_2_, 100 mM KCl pH7.5) was added until a stable baseline is achieved. (6) 1.3 µM of MreB protein in reaction buffer is added for approximately 3-5 min, and the adsorption of MreB was measured for at least 20 min before (7) exchanging and rinsing with reaction buffer.

**Figure Sup 5.**

Low magnification TEM acquisitions reveal remnant of MreB filament structures. MreB (1.3 µM) was set to polymerize in the presence of ATP and a lipid monolayer, before being adsorb on a TEM grid, fixed with uranyl acetate, dried and observed by TEM. **A**. Dual protofilaments are visible, isolated or as tubes or sheets of polymers, at the highest magnification (right-hand panels). Areas of dense concentration of filaments, and particularly bundles and sheets of MreB, can be observed from afar, at lower magnification, due to the differential local increase of uranyl acetate (left). **B.** In limiting amount of ATP (imaged here after 2 to 3h), similar typical dense areas of MreB are still observable at lower magnification suggesting the presence of filaments and sheets (left). Yet, no filaments can be observed at the highest magnification, revealing the local disordering and misalignment of the proteins. The ghost-like image of the ordered structures at low magnification indicates that the local density of proteins was largely conserved, implying a limited motions of the proteins in the timeframe of the experiment.

**S1 Table. Strains used in this study.**

**S2 Table. Plasmids used in this study.**

**S3 Table. Oligonucleotides used in this study.**

**S4 Table. Summary of phenotypes observed in the hereby-studied mutants of MreB**

## References

1. Carballido-Lopez, R. The Actin-like MreB ‘Cytoskeleton’. In: Bacillus: Cellular and Molecular Biology (Third edition) (ed Graumann P. L.). Caister Academic Press (2017).

2. Chastanet, A. & Carballido-Lopez, R. The actin-like MreB proteins in *Bacillus subtilis*: a new turn. Frontiers in bioscience (Scholar edition) 4, 1582–1606 10.2741/S354 (2012).

3. Garner, E. C. Toward a Mechanistic Understanding of Bacterial Rod Shape Formation and Regulation. Annu Rev Cell Dev Biol 37, 1–21 10.1146/annurev-cellbio-010521-010834 (2021).

4. Pompeo, F., et al. RagB stimulates the activity of the peptidoglycan polymerase RodA in *Bacillus subtilis*. EMBO reports 26, 4587–4606 10.1038/s44319-025-00547-w (2025).

5. van den Ent, F., Amos, L. A. & Lowe, J. Prokaryotic origin of the actin cytoskeleton. Nature 413, 39–44 10.1038/35092500 (2001).

6. Ku, C., Lo, W. S., Chen, L. L. & Kuo, C. H. Complete Genome Sequence of Spiroplasma apis B31T (ATCC 33834), a Bacterium Associated with May Disease of Honeybees (Apis mellifera). Genome Announc 2, 10.1128/genomeA.01151-13 (2014).

7. Takahashi, D., Fujiwara, I. & Miyata, M. Phylogenetic origin and sequence features of MreB from the wall-less swimming bacteria Spiroplasma. Biochemical and biophysical research communications 533, 638–644 10.1016/j.bbrc.2020.09.060 (2020).

8. Billaudeau, C., Yao, Z., Cornilleau, C., Carballido-Lopez, R. & Chastanet, A. MreB Forms Subdiffraction Nanofilaments during Active Growth in *Bacillus subtilis*. MBio 10, 10.1128/mBio.01879-18 (2019).

9. Domínguez-Escobar, J., et al. Processive movement of MreB-associated cell wall biosynthetic complexes in bacteria. Science (New York, NY 333, 225–228 10.1126/science.1203466 (2011).

10. Garner, E. C., et al. Coupled, circumferential motions of the cell wall synthesis machinery and MreB filaments in *B. subtilis*. Science (New York, NY 333, 222–225 (2011).

11. Hussain, S., et al. MreB filaments align along greatest principal membrane curvature to orient cell wall synthesis. eLife 7, 10.7554/eLife.32471 (2018).

12. Mao, W., et al. On the role of nucleotides and lipids in the polymerization of the actin homolog MreB from a Gram-positive bacterium. eLife 12, 10.7554/eLife.84505 (2023).

13. van den Ent, F., Izore, T., Bharat, T. A., Johnson, C. M. & Lowe, J. Bacterial actin MreB forms antiparallel double filaments. eLife 3, e02634 10.7554/eLife.02634 (2014).

14. Pande, V., Mitra, N., Bagde, S. R., Srinivasan, R. & Gayathri, P. Filament organization of the bacterial actin MreB is dependent on the nucleotide state. The Journal of cell biology 221, 10.1083/jcb.202106092 (2022).

15. Takahashi, D., Kiyama, H., Matsubayashi, H. T., Fujiwara, I. & Miyata, M. A bacterial actin with high ATPase activity regulates the polymerization of a partner MreB isoform essential for Spiroplasma swimming motility. The Journal of biological chemistry 301, 110462 10.1016/j.jbc.2025.110462 (2025).

16. Wagstaff, J. & Lowe, J. Prokaryotic cytoskeletons: protein filaments organizing small cells. Nature reviews Microbiology 16, 187–201 10.1038/nrmicro.2017.153 (2018).

17. Adriaans, I. E., et al. Intrafilament nucleotide exchange in a prokaryotic actin homolog. Preprint at https://wwwbiorxivorg/content/, 10.64898/2026.07.24.740542 (2026).

18. Salje, J., van den Ent, F., de Boer, P. & Lowe, J. Direct membrane binding by bacterial actin MreB. Molecular cell 43, 478–487 10.1016/j.molcel.2011.07.008 (2011).

19. Nurse, P. & Marians, K. J. Purification and characterization of *Escherichia coli* MreB protein. The Journal of biological chemistry 288, 3469–3475 10.1074/jbc.M112.413708 (2013).

20. Duan, Y., Sperber, A. M. & Herman, J. K. YodL and YisK Possess Shape-Modifying Activities That Are Suppressed by Mutations in *Bacillus subtilis mreB* and *mbl*. Journal of bacteriology 198, 2074–2088 10.1128/JB.00183-16 (2016).

21. Dye, N. A., Pincus, Z., Fisher, I. C., Shapiro, L. & Theriot, J. A. Mutations in the nucleotide binding pocket of MreB can alter cell curvature and polar morphology in *Caulobacter*. Molecular microbiology 81, 368–394 (2011).

22. Ouzounov, N., et al. MreB Orientation Correlates with Cell Diameter in *Escherichia coli*. Biophysical journal 111, 1035–1043 10.1016/j.bpj.2016.07.017 (2016).

23. Gitai, Z., Dye, N. A., Reisenauer, A., Wachi, M. & Shapiro, L. MreB actin-mediated segregation of a specific region of a bacterial chromosome. Cell 120, 329–341 (2005).

24. Shiomi, D., et al. Mutations in cell elongation genes *mreB*, *mrdA* and *mrdB* suppress the shape defect of RodZ-deficient cells. Molecular microbiology 87, 1029–1044 10.1111/mmi.12148 (2013).

25. Yakhnina, A. A. & Gitai, Z. The small protein MbiA interacts with MreB and modulates cell shape in *Caulobacter crescentus*. Molecular microbiology 85, 1090–1104 10.1111/j.1365-2958.2012.08159.x (2012).

26. Maharjan, S., et al. Alanine-scanning mutagenesis library of MreB reveals distinct roles for regulating cell shape and viability. PLoS genetics 22, e1012070 10.1371/journal.pgen.1012070 (2026).

27. Defeu Soufo, H. J. & Graumann, P. L. Dynamic localization and interaction with other *Bacillus subtilis* actin-like proteins are important for the function of MreB. Molecular microbiology 62, 1340–1356 (2006).

28. Billaudeau, C., et al. Contrasting mechanisms of growth in two model rod-shaped bacteria. Nature communications 8, 15370 10.1038/ncomms15370 (2017).

29. Mirouze, N., Ferret, C., Yao, Z., Chastanet, A. & Carballido-Lopez, R. MreB-Dependent inhibition of cell elongation during the escape from competence in *Bacillus subtilis*. PLoS genetics 11, e1005299 10.1371/journal.pgen.1005299 (2015).

30. Jones, L. J., Carballido-López, R. & Errington, J. Control of cell shape in bacteria: helical, actin-like filaments in *Bacillus subtilis*. Cell 104, 913–922 10.1016/S0092-8674(01)00287-2 (2001).

31. Kruse, T., Moller-Jensen, J., Lobner-Olesen, A. & Gerdes, K. Dysfunctional MreB inhibits chromosome segregation in *Escherichia coli*. The EMBO journal 22, 5283–5292 (2003).

32. Adriaans, I. E., et al. ATP-driven membrane binding and polymerization of bacterial actin MreB promotes local membrane fluidization. Biophysical journal, 10.1016/j.bpj.2026.06.003 (2026).

33. De La Cruz, E. M. & Pollard, T. D. Nucleotide-free actin: stabilization by sucrose and nucleotide binding kinetics. Biochemistry 34, 5452–5461 10.1021/bi00016a016 (1995).

34. Kinosian, H. J., Selden, L. A., Estes, J. E. & Gershman, L. C. Nucleotide binding to actin. Cation dependence of nucleotide dissociation and exchange rates. The Journal of biological chemistry 268, 8683–8691 (1993).

35. Dauner, M., Storni, T. & Sauer, U. Bacillus subtilis metabolism and energetics in carbon-limited and excess-carbon chemostat culture. Journal of bacteriology 183, 7308–7317 10.1128/JB.183.24.7308-7317.2001 (2001).

36. Pollard, T. D. Actin and Actin-Binding Proteins. Cold Spring Harbor perspectives in biology 8, 10.1101/cshperspect.a018226 (2016).

37. Huang, W. Z., Wang, J. J., Chen, H. J., Chen, J. T. & Shaw, G. C. The heat-inducible essential response regulator WalR positively regulates transcription of sigI, mreBH and lytE in Bacillus subtilis under heat stress. Research in microbiology 164, 998–1008 10.1016/j.resmic.2013.10.003 (2013).

38. Tseng, C. L. & Shaw, G. C. Genetic evidence for the actin homolog gene mreBH and the bacitracin resistance gene bcrC as targets of the alternative sigma factor SigI of Bacillus subtilis. Journal of bacteriology 190, 1561–1567 (2008).

39. Dobihal, G. S., Brunet, Y. R., Flores-Kim, J. & Rudner, D. Z. Homeostatic control of cell wall hydrolysis by the WalRK two-component signaling pathway in Bacillus subtilis. eLife 8, 10.7554/eLife.52088 (2019).

40. Brunet, Y. R., Habib, C., Brogan, A. P., Artzi, L. & Rudner, D. Z. Intrinsically disordered protein regions are required for cell wall homeostasis in Bacillus subtilis. Genes and development 36, 970–984 10.1101/gad.349895.122 (2022).

41. Dominguez-Cuevas, P., Porcelli, I., Daniel, R. A. & Errington, J. Differentiated roles for MreB-actin isologues and autolytic enzymes in Bacillus subtilis morphogenesis. Molecular microbiology 89, 1084–1098 10.1111/mmi.12335 (2013).

42. Liu, Y., et al. Membrane binding properties of the cytoskeletal protein bactofilin. eLife 13, 10.7554/eLife.100749 (2025).

43. Harwood, C. R. & Cutting, S. M. Molecular Biological Methods for Bacillus. (J. Wileys and Sons, New York, 1990).

44. Gibson, D. G., et al. Enzymatic assembly of DNA molecules up to several hundred kilobases. Nature methods 6, 343–345 10.1038/nmeth.1318 (2009).

45. Vellanoweth, R. L. & Rabinowitz, J. C. The influence of ribosome-binding-site elements on translational efficiency in Bacillus subtilis and Escherichia coli in vivo. Molecular microbiology 6, 1105–1114 (1992).

46. Landgraf, D., Okumus, B., Chien, P., Baker, T. A. & Paulsson, J. Segregation of molecules at cell division reveals native protein localization. Nature methods 9, 480–482 10.1038/nmeth.1955 (2012).

47. Cornilleau, C., Chastanet, A., Billaudeau, C. & Carballido-Lopez, R. Methods for Studying Membrane-Associated Bacterial Cytoskeleton Proteins In Vivo by TIRF Microscopy. Methods Mol Biol 2101, 123–133 10.1007/978-1-0716-0219-5_8 (2020).

48. Ducret, A., Quardokus, E. M. & Brun, Y. V. MicrobeJ, a tool for high throughput bacterial cell detection and quantitative analysis. Nature microbiology 1, 10.1038/nmicrobiol.2016.77 (2016).

49. Schindelin, J., et al. Fiji: an open-source platform for biological-image analysis. Nature methods 9, 676-682 10.1038/nmeth.2019 (2012).

50. Billaudeau, C., Chastanet, A. & Carballido-Lopez, R. Processing TIRF Microscopy Images to Characterize the Dynamics and Morphology of Bacterial Actin-Like Assemblies. Methods Mol Biol 2101, 135–145 10.1007/978-1-0716-0219-5_9 (2020).

51. Ferhan, A. R., et al. Solvent-assisted preparation of supported lipid bilayers. Nat Protoc 14, 2091–2118 10.1038/s41596-019-0174-2 (2019).

52. Rodahl, M., Hook, F., Krozer, A., Brzezinski, P. & Kasemo, B. Quartz-Crystal Microbalance Setup for Frequency and Q-Factor Measurements in Gaseous and Liquid Environments. Rev Sci Instrum 66, 3924–3930 10.1063/1.1145396 (1995).

53. Renner, L. D. & Weibel, D. B. MinD and MinE interact with anionic phospholipids and regulate division plane formation in *Escherichia coli*. The Journal of biological chemistry 287, 38835–38844 10.1074/jbc.M112.407817 (2012).

