## Supplementary materials tables and figures for "A genetic screen unveils key molecular steps in the polymerization cycle of the bacterial actin-like MreB"

de San Eustaquio *et al*. Figure Sup 1

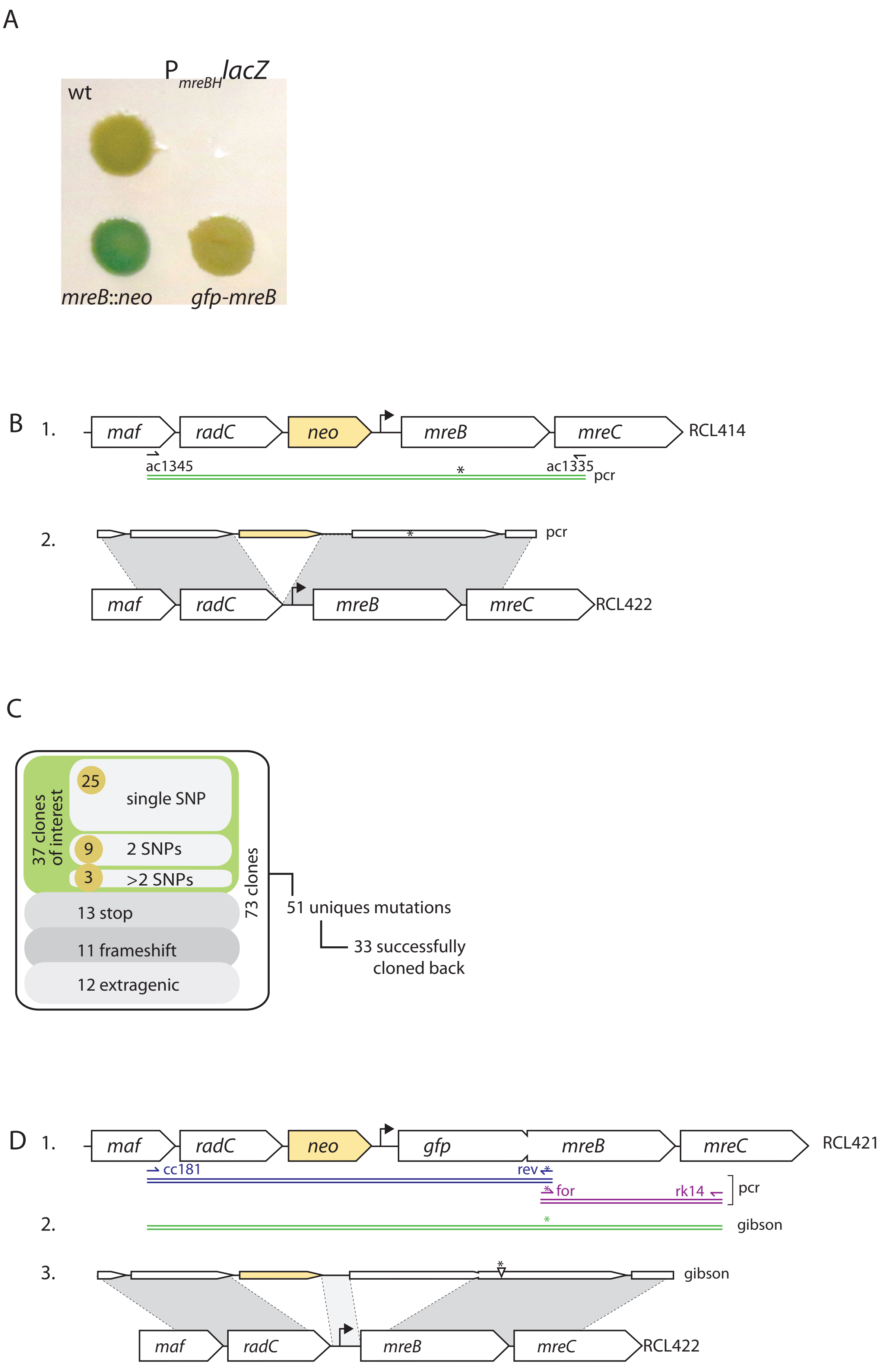

E

```
              β1      β2       β3        β4           β5   α1
T.mar   -------MLRKDIGIDLGTANTLVFLRGKGIVVNEPSVIAIDST----TGEILKVGLEAK  49
B.sub   ----MFGIGARDLGIDLGTANTLVFVKGKGIVVREPSVVALQTD----TKSIVAVGNDAK  52
G.ste   ----MFGIGTKDLGIDLGTANTLVYVKGKGIVLREPSVVAIQRD----TKQIVAVGNEAK  52
E.col   MLKKFRGMFSNDLSIDLGTANTLIYVKGQGIVLNEPSVVAIRQDRAGSPKSVAAVGHDAK  60
C.cre   MFSSLFGVISNDIAIDLGTANTLIYQKGKGIVLNEPSVVALRN--VGGRKVVHAVGIEAK  58
              :  .*:.*********::  :*:***:.****:*:          :  **  :**
                        14

        α1    β6                α2                    β7
T.mar   NMIGKTPATIKAIRPMRDGVIADYTVALVMLRYFINKAKGGMNLF--KPRVVIGVPIGIT 107
B.sub   NMIGRTPGNVVALRPMKDGVIADYETTATMMKYYINQAIKNKGMFTRKPYVMVCVPSGIT 112
G.ste   NMIGRTPGNIVALRPMKDGVIADYETTATMMKYYIRKAIKTKGLFAGKPYVMVCVPYGIT 112
E.col   QMLGRTPGNIAAIRPMKDGVIADFFVTEKMLQHFIKQVHSNSFMRP-SPRVLVCVPVGAT 119
C.cre   QMLGRTPGHMEAIRPMRDGVIADFEVAEEMIKYFIRKVHNRKG-FV-NPKVIVCVPSGAT 116
        :  :*:**. :  *:** :******: .:    ::::*.:.      .*  *:: **  *  *
          56

          α3              β8    α4         β9        β10
T.mar   DVERRAILDAGLEAGASKVFLIEEPMAAAIGSNLNVEEPSGNMVVDIGGGTTEVAVISLG 167
B.sub   AVEERAVIDATRQAGARDAYPIEEPFAAAIGANLPVWEPTGSMVVDIGGGTTEVAIISLG 172
G.ste   AVEERAVIDATRQAGARDAYTIEEPFAAAIGANLPVWEPTGSMVVDIGGGTTEVAVISLG 172
E.col   QVERRAIRESAQGAGAREVFLIEEPMAAAIGAGLPVSEATGSMVVDIGGGTTEVAVISLN 179
C.cre   AVERRAINDSCLNAGARRVGLIDEPMAAAIGAGLPIHEPTGSMVVDIGGGTTEVAVLSLS 176
         **.**: ::   ***   .  *:** *****:.* : * :*. ************::**.
         114                    136                   158 160       171

        β11          α5            β12   α6                β13
T.mar   SIVTWESIRIAGDEMDEAIVQYVRETYRVAIGERTAERVKIEIGNVFPSKENDELETTVS 227
B.sub   GIVTSQSIRVAGDEMDDAIINYIRKTYNLMIGDRTAEAIKMEIGSAEAP--EESDNMEIR 230
G.ste   GIVTSQSIRIAGDEMDEAIIQYIRKSYNLMIGERTAEAIKMEIGSAGNP--EGIGNMEIR 230
E.col   GVVYSSSVRIGGDRFDEAIINYVRRNYGSLIGEATAERIKHEIGSAYPG--DEVREIEVR 237
C.cre   GIVYSRSVRVGGDKMDEAIISYMRRHHNLLIGETTAERIKKEIGTARAPADGEGLSIDVK 236
        .:*   *:*:*:.**. *:**:.*:*. :   **:  *** :*  ***..          .  :

        β13     β14      α7                       α8     β15
T.mar   GIDLSTGLPRKLTLKGGEVREALRSVVVAIVESVRTTLEKTPPELVSDIIERGIFLTGGG 287
B.sub   GRDLLTGLPKTIEITGKEISNALRDTVSTIVEAVKSTLEKTPPELAADIMDRGIVLTGGG 290
G.ste   GRDLLTGLPKTIEISAEEVAEALRDTVYAIVESVKNTLEKTPPELAADIMDRGIVLTGGG 290
E.col   GRNLAEGVPRGFTLNSNEILEALQEPLTGIVSAVMVALEQCPPELASDISERGMVLTGGG 297
C.cre   GRDLMQGVPREVRISEKQAADALAEPVGQIVEAVKVALEATPPELASDIADKGIMLTGGG 296
        * :*   *:*:  . :.   :  :**  . :  **.:*   :**  ****.:**  ::*:.*****
        231

                        β16      α10      α11
T.mar   SLLRGLDTLLQKETGISVIRSEEPLTAVAKGAGMVLDKVNILKKLQGAGGS     338
B.sub   ALLRNLDKVISEETKMPVLIAEDPLDCVAIGTGKALEHIHLFKGKTR----     337
G.ste   ALLRNLDKVISQETDMPVIVAENPLDCVAIGTGKALDHIDLFKNKARDHR-     340
E.col   ALLRNLDRLLMEETGIPVVVAEDPLTCVARGGGKALEMIDMHGGDLFSEE-     347
C.cre   ALLRGLDAEIRDHTGLPVTVADDPLSCVALGCGKVLEHPKWMKGVLESTLA     347
        :***.**   :  ..* : *   :::** .** * *  .*:   .
```

Fig. Sup2

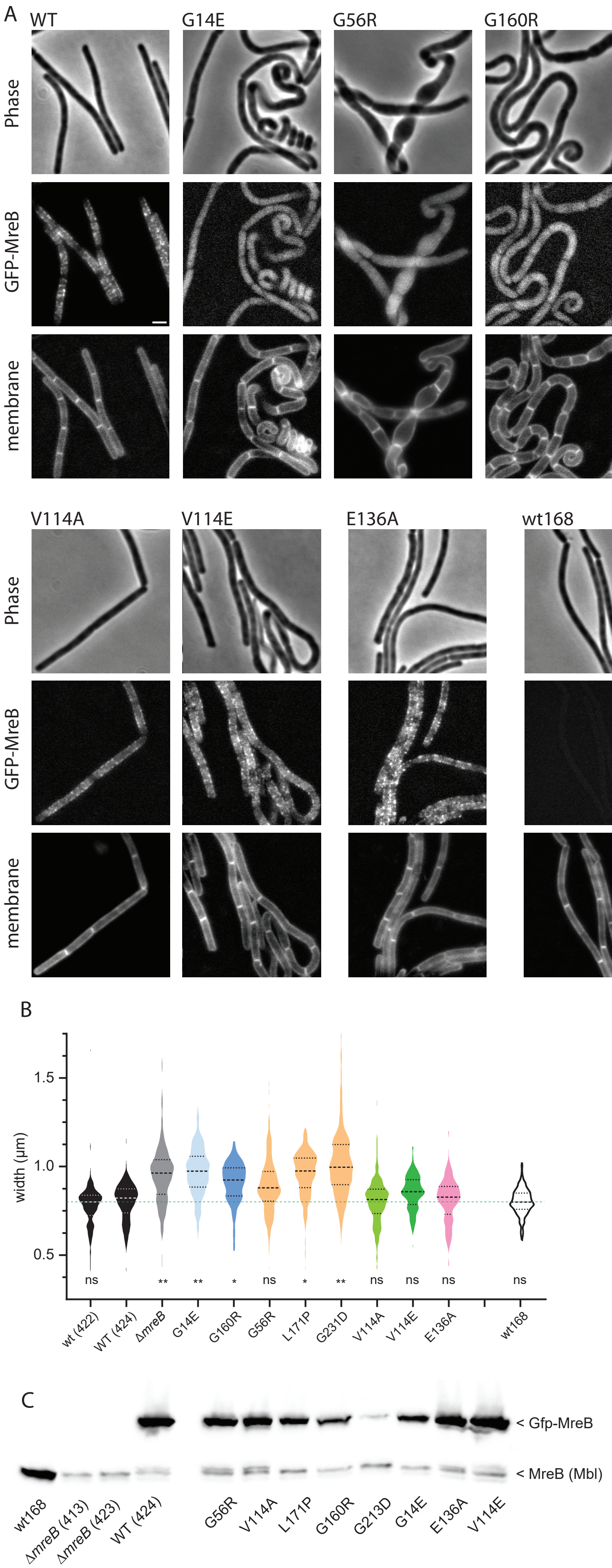
A
WT
G14E
G56R
G160R
L171P
G231D
Phase
GFP-MreB
membrane
V114A
V114E
E136A
wt168
ΔmreB
B
width (µm)
1.5
1.0
0.5
ns
**
**
*
ns
*
**
ns
ns
ns
ns
wt (422)
WT (424)
ΔmreB
G14E
G160R
G56R
L171P
G231D
V114A
V114E
E136A
wt168
C
< Gfp-MreB
< MreB (Mbl)
wt168
ΔmreB (413)
ΔmreB (423)
WT (424)
G56R
V114A
L171P
G160R
G213D
G14E
E136A
V114E

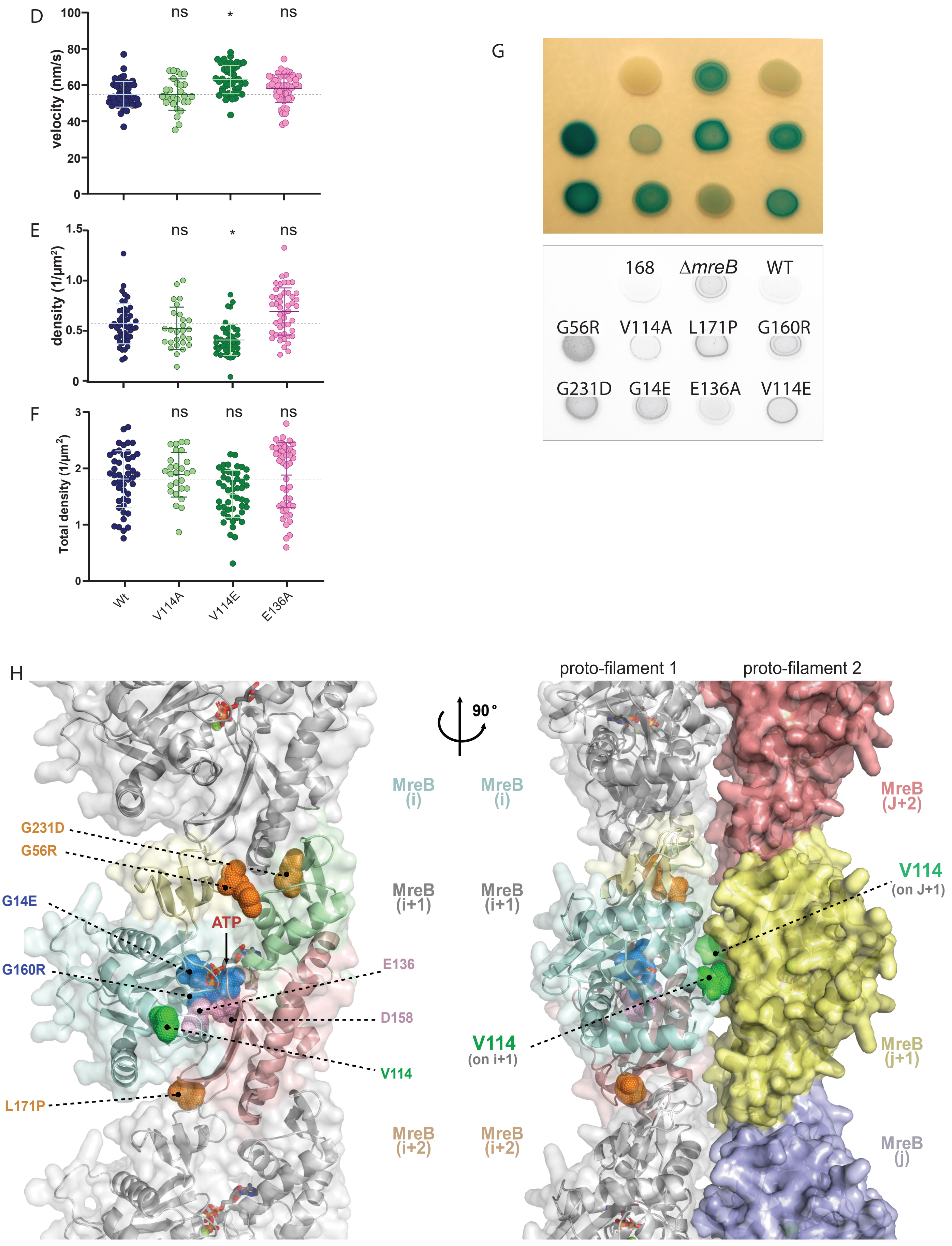
D
velocity (nm/s)
100
80
60
40
20
0
ns
*
ns
E
density (1/µm²)
1.5
1.0
0.5
0
ns
*
ns
F
Total density (1/µm²)
3
2
1
0
ns
ns
ns
Wt
V114A
V114E
E136A
G
168
ΔmreB
WT
G56R
V114A
L171P
G160R
G231D
G14E
E136A
V114E
H
proto-filament 1
proto-filament 2
90°
MreB (i)
MreB (i+1)
MreB (i+2)
MreB (J+2)
MreB (j+1)
MreB (j)
G231D
G56R
G14E
ATP
G160R
E136
D158
V114
L171P
V114 (on J+1)
V114 (on i+1)

de San Eustaquio *et al*. Figure Sup 3

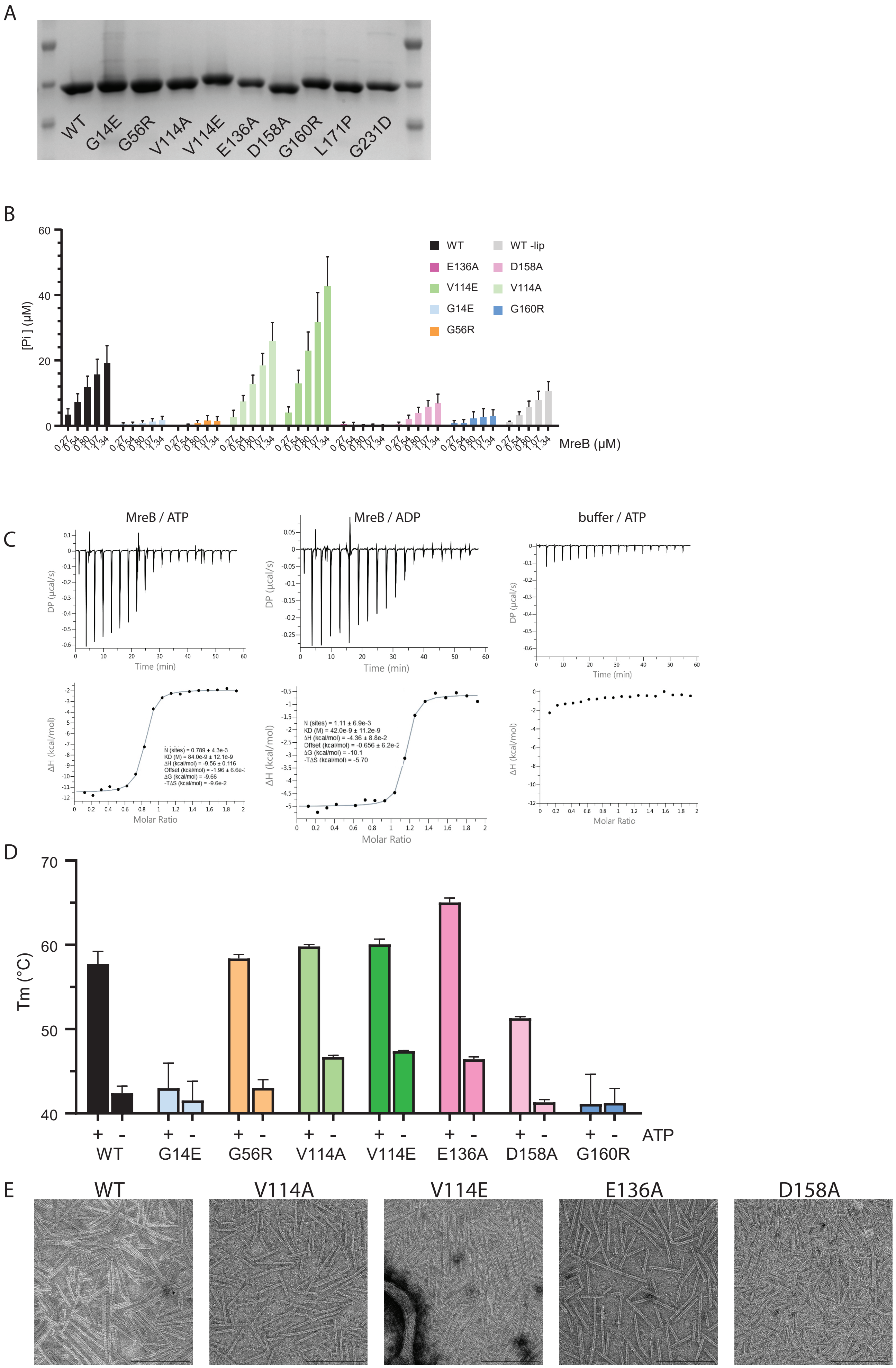

Fig 4 sup

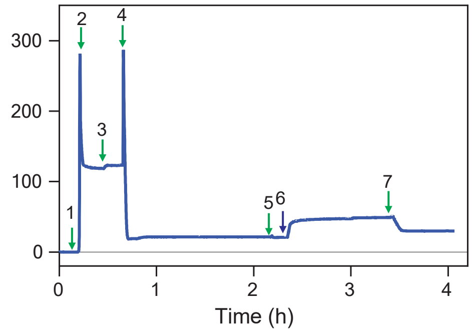

Fig. Sup. 5

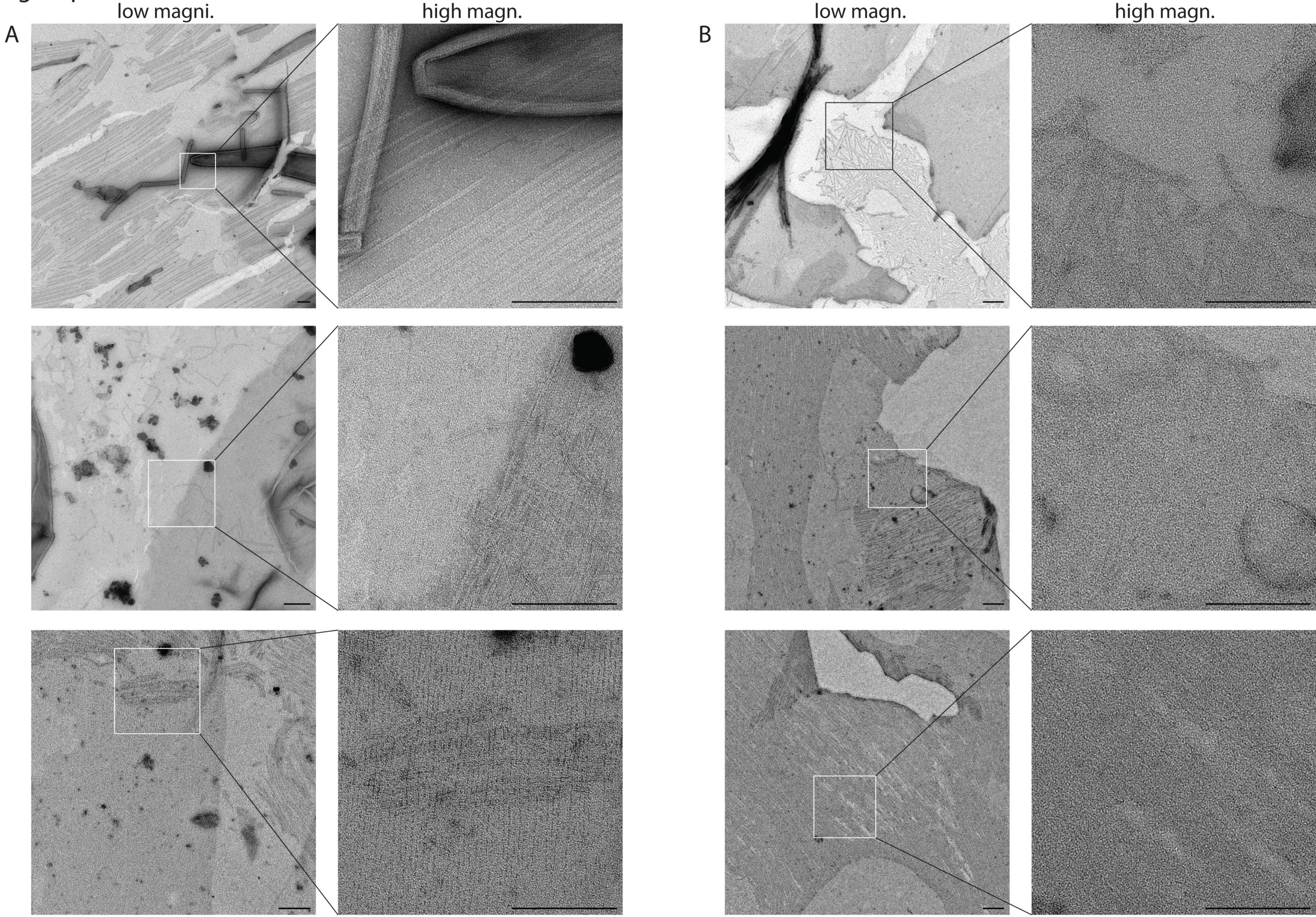

#### Sup. Table 1. Strains used in this study

| Strain | Relevant genotype [1] | Source or reference [2] |
|---|---|---|
| *B. subtilis* | | |
| 168 | *trpC2,* (wt) | Laboratory stock |
| RCL413 | *trpC2,* Δ*mreB* Ω*neo* | Billaudeau *et al.,* 2019 |
| RCL414 | *trpC2, mreB* Ω*neo* | OEPCR → 168 (see M&M) |
| RCL238 | *trpC2,* Δ*mreB* Ω*neo, amyE* ::($P_{xyl}$ *gfp-mreB, spc)* | Billaudeau *et al.* , 2017 |
| ABS1755 | *trpC2, thrC* ::($P_{mreBH}$ *lacZ, spc* ) | pAC783 → 168 (see M&M) |
| RCL422 | *trpC2, thrC* ::($P_{mreBH}$ *lacZ, spc ::cat* ) | pDag32 → ABS1755 (see M&M) |
| RCL423 | *trpC2,* Δ*mreB* Ω*neo, thrC* ::($P_{mreBH}$ *lacZ, spc::cat* ) | RCL413 → RCL422 |
| RCL420 | *trpC2 , mreB* Ω(*neo* , $P_{nat (o.r)}$ *gfp-mreB* ) | ITA → 168 (see M&M) |
| RCL421 | *trpc2, mreB* Ω(*neo* $P_{nat (o.r)}$ *gfp* $^{A206K}$ *mreB* ) | ITA → 168 (see M&M) |
| RCL424 | *trpC2, thrC* ::($P_{mreBH}$ *lacZ, spc::cat* ), *mreB* Ω(*neo,* $P_{nat (o.r)}$ *gfp* $^{A206K}$ *mreB* ) | RCL421 → RCL422 |
| RCL0427 | *trpC2, thrC* ::($P_{mreBH}$ *lacZ, spc::cat* ), *mreB* ::(*neo-* $P_{nat (o.r)}$ *gfp* $^{A206K}$ *mreB* $^{G56R}$ ) | ITA → RCL422 |
| RCL0429 | *trpC2, thrC* ::($P_{mreBH}$ *lacZ, spc::cat* ), *mreB* ::(*neo-* $P_{nat (o.r)}$ *gfp* $^{A206K}$ *mreB* $^{G160R}$ ) | ITA → RCL422 |
| RCL1212 | *trpC2, thrC* ::($P_{mreBH}$ *lacZ, spc::cat* ), *mreB* ::(*neo-* $P_{nat (o.r)}$ *gfp* $^{A206K}$ *mreB* $^{G160R}$ ) | RCL429 → RCL422 |
| RCL0433 | *trpC2, thrC* ::($P_{mreBH}$ *lacZ, spc::cat* ), *mreB* ::(*neo-* $P_{nat (o.r)}$ *gfp* $^{A206K}$ *mreB* $^{V114A}$ ) | ITA → RCL422 |
| RCL1255 | *trpC2, thrC* ::($P_{mreBH}$ *lacZ, spc::cat* ), *mreB* ::(*neo-* $P_{nat (o.r)}$ *gfp* $^{A206K}$ *mreB* $^{V114E}$ ) | ITA → RCL422 |
| RCL0438 | *trpC2, thrC* ::($P_{mreBH}$ *lacZ, spc::cat* ), *mreB* ::(*neo-* $P_{nat (o.r)}$ *gfp* $^{A206K}$ *mreB* $^{G231D}$ ) | ITA → RCL422 |
| RCL1213 | *trpC2, thrC* ::($P_{mreBH}$ *lacZ, spc::cat* ), *mreB* ::(*neo-* $P_{nat (o.r)}$ *gfp* $^{A206K}$ *mreB* $^{G231D}$ ) | RCL438 → RCL422 |
| RCL0441 | *trpC2, thrC* ::($P_{mreBH}$ *lacZ, spc::cat* ), *mreB* ::(*neo-* $P_{nat (o.r)}$ *gfp* $^{A206K}$ *mreB* $^{G14E}$ ) | ITA → RCL422 |
| RCL1214 | *trpC2, thrC* ::($P_{mreBH}$ *lacZ, spc::cat* ), *mreB* ::(*neo-* $P_{nat (o.r)}$ *gfp* $^{A206K}$ *mreB* $^{G14E}$ ) | RCL441→ RCL422 |
| RCL0449 | *trpC2, thrC* ::($P_{mreBH}$ *lacZ, spc::cat* ), *mreB* ::(*neo-* $P_{nat (o.r)}$ *gfp* $^{A206K}$ *mreB* $^{L171P}$ ) | ITA → RCL422 |
| RCL1209 | *trpC2, thrC* ::($P_{mreBH}$ *lacZ, spc::cat* ), *mreB* ::(*neo-* $P_{nat (o.r)}$ *gfp* $^{A206K}$ *mreB* $^{L171P}$ ) | RCL449-> RCL422 |
| RCL1254 | *trpC2, thrC* ::($P_{mreBH}$ *lacZ, spc::cat* ), *mreB* ::(*neo-* $P_{nat (o.r)}$ *gfp* $^{A206K}$ *mreB* $^{E136A}$ ) | ITA → RCL422 |
| *E. coli* | | |
| T7 express [3] | *F- λ- fhuA2* [*lon* ] *ompT lacZ* ::*T7 gene1 gal sulA11* Δ(*mcrC-mrr* )*114* ::*IS10 R* (*mcr-73* ::*miniTn10-TetS* )*2 R(zgb-210* ::*Tn10* )(*TetS* ) *endA1* [*dcm* ] | New England Biolabs |
| EcRCL212 | p ($P_{T7}$ *6his-mreB* $_{Gst}$ ) | pCC110 -> T7 express |
| EcRCL347 | p ($P_{T7}$ *6his-mreB* $_{Gst}^{G14E}$ ) | pLH2 → T7 express |
| EcRCL547 | p ($P_{T7}$ *6his-mreB* $_{Gst}^{G56R}$ ) | pSA3 → T7 express |
| EcRCL556 | p ($P_{T7}$ *6his-mreB* $_{Gst}^{V114A}$ ) | pSA7 → T7 express |
| EcRCL550 | p ($P_{T7}$ *6his-mreB* $_{Gst}^{V114E}$ ) | pSA6 → T7 express |
| EcRCL545 | p ($P_{T7}$ *6his-mreB* $_{Gst}^{E136A}$ ) | pSA1 → T7 express |
| EcRCL551 | p ($P_{T7}$ *6his-mreB* $_{Gst}^{D158A}$ ) | pSA11 → T7 express |
| EcRCL349 | p ($P_{T7}$ *6his-mreB* $_{Gst}^{G160R}$ ) | pLH4 → T7 express |
| EcRCL548 | p ($P_{T7}$ *6his-mreB* $_{Gst}^{L171P}$ ) | pSA4 → T7 express |
| EcRCL617 | p ($P_{T7}$ *6his-mreB* $_{Gst}^{G231D}$ ) | pCM003 → T7 express |

[1] p(xxx) stand for plasmids
[2] Arrows indicate transformation direction; ITA stands for isothermal assembly; OEPCR for overlapping extension PCR
[3] expression strain carrying the T7 RNA polymerase gene under *lac* promoter control, allowing IPTG-induced expression of a gene of interest

#### Sup. Table 2. Plasmids used in this study

| name | Relevant genotype | Source or reference[1] |
|---|---|---|
| pDG1729 | p (*thrC* ::(*lacZ, spc* ), *mls* , *bla* ) | Guerout-Fleury *et al.,* 1996[4] |
| pAC783 | p (*thrC* ::($P_{mreBH}$ *lacZ, spc* ), *mls* , *bla* ) | pDG1729 derivative |
| pDag32 | p (Δ*spc* ::*cat, bla* ) | laboratory stock[2] |
| pCC110 | p ($P_{T7}$ *6his-mreB* $_{Gst}$ , *km* ) | Mao *et al* ., 2023[3] |
| pLH2 | p ($P_{T7}$ *6his-mreB* $_{Bsu}^{G14E}$ ) | QC on pCC110 with ac1395/1396 |
| pSA3 | p ($P_{T7}$ *6his-mreB* $_{Bsu}^{G56R}$ ) | QC on pCC110 with ac1413/ac1414 |
| pSA7 | p ($P_{T7}$ *6his-mreB* $_{Bsu}^{V114A}$ ) | QC on pCC110 with ac1421/ac1422 |
| pSA6 | p ($P_{T7}$ *6his-mreB* $_{Bsu}^{V114E}$ ) | QC on pCC110 with ac1417/ac1418 |
| pSA1 | p ($P_{T7}$ *6his-mreB* $_{Bsu}^{E136A}$ ) | QC on pCC110 with ac1407/ac1408 |
| pSA11 | p ($P_{T7}$ *6his-mreB* $_{Bsu}^{D158A}$ ) | QC on pCC110 with ac1423/ac1410 |
| pLH4 | p ($P_{T7}$ *6his-mreB* $_{Bsu}^{G160R}$ ) | QC on pCC110 with ac1397/ac1398 |
| pSA4 | p ($P_{T7}$ *6his-mreB* $_{Bsu}^{L171P}$ ) | QC on pCC110 with ac1415/ac1416 |
| pCM3 | p ($P_{T7}$ *6his-mreB* $_{Bsu}^{G231D}$ ) | QC on pCC110 with asec179/asec177 |

[1] QC stands for quickchange; see M&M for details
[2] From the Losick laboratory stock; similar to that discussed in : Steinmetz & Richter, Gene, 142 (1994) 79-83.
[3] DOI: 10.7554/eLife.84505
[4] DOI: 10.1016/S0378-1119(96)00404-0

#### Sup. Table 3. Oligonucleotides used in this study

| name | sequence | use |
|---|---|---|
| ac1345 | AAGTCACTCAGTAATAACCGC | RCL414 (neo at mreB locus) |
| ac1346 | GAAAAACTCATAAACTCTATTATAGC | " |
| ac1334 | TTTTTTTCGTCGAATTAAGCTATAATAGA | " |
| ac1335 | TCGATCAAGCCGTAGCCTTTGCTG | " |
| ac1246 | GATGAATTCTTCATCTACTTTTCTCACAACA | ABS1755 ($P_{mreBH}$ *lacZ* ) |
| ac1247 | GGAGGATCCCCTAATTTAATATGATTCTACATTT | " |
| cc181 | GTCATGGGCCTTCCTATATC | RCL421 ($P_{nat}$ *gfpmeB* ) |
| cc186 | TAGTTTACCTCCTTAAATGTATGTATCTTCCTTTCTTAAAGC | " |
| cc182 | GGATGTGCTCCAGTGCTTTC | " |
| cc185 | GATACATACATTTAAGGAGGTAAACTAATGAGTAAAGGAGAAGAACTTTTCACTG | " |
| ac1281 | CAATCTAAACTTTCGAAAGATCCCAACG | quickchange on *gfp* (A206K) |
| ac1282 | CGAAAGTTTAGATTGTGTGGACAGGTAATG | " |
| RK14 | AATTCGAGCAGACAGACAGCCAGAAC | |
| asec91 | GCCCGGTGTCCGTC<u>T</u>AATCATATTTTTCGC | QC for expression in *B. subtilis* of G56R |
| asec92 | ATATGATT<u>A</u>GACGGACACCGGGCAACGTGG | " |
| asec95 | GTACCGCCC<u>C</u>TGATATCAACAACCATGCTTCC | QC for expression in *B. subtilis* of G160R |
| asec96 | TGTTGATATCA<u>G</u>GGGCGGTACGACAGAAGTTGC | " |
| asec154 | CAGCTGCTGAAGAACGCGCTGTTATCGATGCGACAAGACAGG | QC for expression in *B. subtilis* of V114A |
| asec155 | CGCGTTCTTCAACAGCTGTAATGCCTGATGGG | " |
| asec176 | GAAATCCGCGACCGCGATTTGCTCACAGGTTTGC | QC for expression in *B. subtilis* of G231D |
| asec177 | CGCGGTCGCGGATTTCCATGTTG | " |
| asec182 | ATAGATCTTGAAACTGCGAATACGCTTGTTTTTGT | QC for expression in *B. subtilis* of G14E |
| asec183 | TTCGCAGTTTCAAGATCTATACCAAGGTCTC | " |
| asec229 | TTATTTCCCCCGGAGGCATCGTAACGTC | QC for expression in *B. subtilis* of L171P |
| asec230 | GATGCCTCCGGGGGAAATAATC | " |
| cc270 | ACAGCTGAGGAAGAACGCGCTGTTATC | QC for expression in *B. subtilis* of V114E |
| cc271 | GTTCTTCCTCAGCTGTAATGCCT | " |
| cc272 | ATTGAAGCGCCTTTTGCCGCAGC | QC for expression in *B. subtilis* of E136A |
| cc273 | GCAAAAGGCGCTTCAATCGGATACG | " |
| | <u>Primers for construction of plasmids for heterologous expression in *E. coli*</u> | |
| ac1395 | TTTAGAAACAGCGAACACGCTTGTTTAT | QC for expression in *E. coli* of G14E |
| ac1396 | GTGTTCGCTGTTTCTAAATCGATCCCAAGATCTTTCGTTCC | " |
| ac1413 | ATCCGCCGCACGCCCGGCAA | QC for expression in *E. coli* of G56R |
| ac1414 | GTGCGGCGGATCATGTTTTTCGCCTC | " |
| ac1421 | ACAGCGGCGGAGGAGCGGG | QC for expression in *E. coli* of V114A |
| ac1422 | TCCTCCGCCGCTGTAATGCCATA | " |
| ac1417 | ACAGCGGAAGAGGAGCGGG | QC for expression in *E. coli* of V114E |
| ac1418 | TCCTCTTCCGCTGTAATGCCATA | " |
| ac1407 | ACGATTGAAGCGCCGTTTGCGGCTG | QC for expression in *E. coli* of E136A |
| ac1408 | CGCAAACGGCGCTTCAATCGTATAGGCG | " |
| ac1423 | GTCGTCGCGATCGGCGGC | QC for expression in *E. coli* of D158A |
| ac1410 | GCCGCCGATCGCGACGACCATGCTG | " |
| ac1397 | GACATCCGTGGCGGCACGACCGAAGTGGCGGTCA | QC for expression in *E. coli* of G160R |
| ac1398 | CGTGCCGCCACGGATGTCGACGACCATGCTGCCAGT | " |
| ac1415 | GTCATTTCGCCGGGCGGCATCGT | QC for expression in *E. coli* of L171P |
| ac1416 | GCCCGGCGAAATGACCGCCACTT | " |
| asec179 | GTTGTGAGAGGGCCGTCAGTTGTCGCTTTGCAG | QC for expression in *E. coli* of G231D |
| asec177 | CGCGGTCGCGGATTTCCATGTTG | " |

QC stands for quickchange mutagenesis

#### Sup. Table 4. Summary of phenotypes observed in the hereby studied mutants of MreB

| Mutation | | *in vivo* Phenotypes [1] | | | | | *in vitro* Phenotypes [2] | | | | | |
|---|---|---|---|---|---|---|---|---|---|---|---|---|
| Change | Localization[3] | Cell shape (Fig. 2A,C, Sup2A,B) | MreB local. (Fig. 2A, Sup2A) | Foci dynamics (Fig. Sup2D-F) | $P_{mreBH}$ induc. (Fig. Sup2G) | Protein levels (Fig. 2D, Sup2C) | Protein Folding (Fig. 3A) | Polymerization (Fig. 3B, Sup3E) | Pi release (Fig. 3C, Sup3B) | ATP binding (Fig. 3E, Sup3D) | Lipid binding (Fig. 4B, Sup4B) | Polym. stability (Fig. 5) |
| WT | | ref. | ref. | ref. | ref. | ref. | ref. | ref. | ref. | ref. | +++ | ref. |
| L171P | Longit. interf. | = Δ*mreB* | cytosol, diffuse | nd | = Δ*mreB* | intermed | - | - | nd | nd | nd | nd |
| G231D | Longit. interf. | = Δ*mreB* | cytosol, diffuse | nd | > Δ*mreB* | lowest | - | - | nd | nd | nd | nd |
| G56R | Longit. interf. | = Δ*mreB* | cytosol, diffuse | nd | > Δ*mreB* | wt | WT | - | ↓12x | WT | - | nd |
| G14E | catalytic cleft | = Δ*mreB* | cytosol, diffuse | nd | = Δ*mreB* | intermed | WT | - | ↓12x | - | - | nd |
| G160R | catalytic cleft | = Δ*mreB* | cytosol, diffuse | nd | = Δ*mreB* | intermed | WT | - | ↓ 6x | - | - | nd |
| D158A | catalytic cleft | nd | nd | nd | nd | nd | WT | WT | ↓ 3x | intermed | nd | nd |
| E136A | catalytic cleft | wt | wt | ≈wt | intermed | wt | WT | WT | ↓55x | > WT | ++ | >> WT |
| V114A | Lateral interf. | wt | wt | ≈wt | intermed | wt | WT | WT | WT (↑ 1,2x) | WT | ++ | nd |
| V114E | Lateral interf. | wt | wt | ↑speed,↓density | = Δ*mreB* | wt | WT | WT | ↑ 2x | WT | +++ | nd |

[1] Phenotypes assassed according to: Phase contrast microscopy imaging (Cell shape), subcellular localization of GFP-MreB (MreB local.), MSD analysis of TIRF imaging (Foci dynamics), induction of the reporter fusion ($P_{mreBH}$ induc.), Western Blotting (MreB levels)
[2] Phenotypes assassed according to: Circular Dichroism (Protein folding), sqTEM (Polymerization), Malachite green assay (Pi release), Thermal shift assay (ATP binding), QCM-D (lipid binding), sqTEM under limiting ATP (polymer stability)
[3] longit. interf. stands for longitudinal intra-protofilament interfaces. Lateral interf. for interprotofilament lateral interface
nd: not determined

### Structural mapping of MreB–MreB and nucleotide-binding interfaces and predicted effects of the analyzed mutations.

**(A)** Structure-based analysis of lateral and longitudinal MreB–MreB interfaces.

**(B)** Predicted effects of the *G. stearothermophilus* MreB mutations on the MreB–MreB interfaces.

**(C)** Nucleotide-interacting residues.

**(D)** Mapping of the analyzed *G. stearothermophilus* MreB mutations relative to these nucleotide-interacting residues. Most identified interactions are shown in Fig Sup. 1E; only a subset is omitted for clarity.

---

**(A) Structure-based analysis of MreB–MreB interfaces.** Lateral and longitudinal protofilament interfaces were analyzed in the crystal structure of AMPPNP–$Mg^{2+}$-bound *C. crescentus* MreB (PDB 4CZJ), which contains two antiparallel protofilaments in the crystal lattice (van den Ent et al., 2014). Interface residues for which ≥40% of the solvent-accessible surface area (SASA) becomes buried upon interface formation were identified using PISA (Krissinel and Henrick, 2007).

**Notation:** * indicates an intermolecular salt bridge with a maximum side-chain interatomic distance of ≤4.0 Å; (+), residues with approximately 90–100% buried SASA; (−), residues near the 40% inclusion threshold.

**lateral (inter-protofilament) interfaces** between two adjacent MreB subunits from neighboring protofilaments. Below are highly buried residues in subdomains:

**Large** (total buried SASA: 1,802 Å²).

| | **I** | **II** |
|---|---|---|
| **B** | none | S197(−) R200*<br>(E275)(−)<br>R201*(E268, E275) |
| **A** | I79 A80 F82<br>T116 A117<br>V118(+) E119<br>R121 A122(+) | R185 E261<br>E268*(R201)<br>K271(−)<br>E275*(R200, R201) |

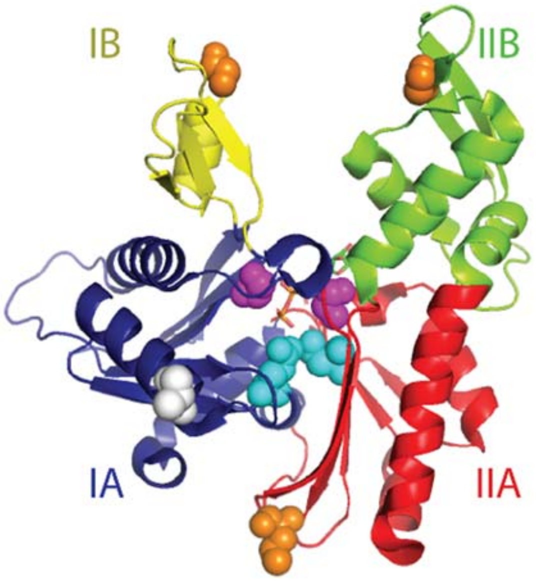


**Small** (total buried SASA: 495 Å²).

| | | |
|---|---|---|
| **B** | none | R200<br>N204(−)<br>L240 |
| **A** | none | none |

**Large longitudinal (intra-protofilament) interface** between adjacent subunits within the same protofilament (total buried SASA: 1,744 Å$^2$)**I**

| | I | II |
|---|---|---|
| **B** | M60(+) G62(+) R63*(E280, D284) T64(−) P65(+) H67(−) M68 | L206(−) I207 G208(+) T211 K236*(D287) G237(+) R238 G243(+) V244(+) P245(+) |
| **A** | G62(+), R63*(E280, D284) T64(−) P65(+) H67(−) | P155(−) S174(+) L175(+) G177(+) I178(−) V179 K271(−) L274(+) E275 T277(+) P279(+) E280*(R63) L281(+) A282(+) S283 D284*(R63) A286 D287*(K236) L312(−) |

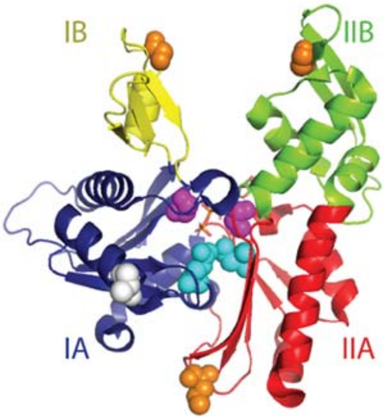


**(B) Proximity mapping of the *G. stearothermophilus* MreB mutations mapped into *C. crescentus* MreB (PDB 4CZJ) and prediction of interface residues.** The mutations were mapped onto the corresponding *C. crescentus* residues. Interface residues with ≥40% buried SASA and located within ±4 positions in the aligned sequences are reported to indicate which interface each mutation may perturb. The UniProt identifiers for *C. crescentus* and *G. stearothermophilus* MreB are A0A0H3C7V4 and A0A150MJ77, respectively; the proteins comprise 347 and 340 residues.

| **Mutated residue in *G. stearothermophilus* MreB** | | | | **Nearby interface residue(s) (±4 residues) in *C. crescentus* MreB** | | **Structure-based interpretation: mutation likely to primarily perturb** |
|---|---|---|---|---|---|---|
| **mutation** | **structural location** | **sub domain** | **in *Cc**** | **longitudinal** (lies near:) | **lateral** (lies near:) | |
| **G14E** | **ATP-binding site** | IA | G18 | None | None | ATP/ADP binding, ATP hydrolysis |
| **G160R** | | IIA | G164 | None | None | ATP/ADP binding, ATP hydrolysis |
| **E136A** | **ATP-binding site** | IA | E140 | None | None | ATP-hydrolysis |
| **D158A** | | IIA | D162 | None | None | ATP/ADP-Mg2+-binding, ATP-hydrolysis |
| **G56R** | **Longitudinal proto-filament interface** | IB | G62 | M60(+), G62(+), R63*(E280, D284), T64(−), P65(+) | None | longitudinal MreB–MreB contacts |
| **L171P** | | IIA | L175 | S174(+), L175(+), G177(+), I178(−), V179 | None | longitudinal MreB–MreB contacts |
| **G231D** | | IIB | G237 | K236*(D287), G237(+), R238 | L240, M241 | longitudinal MreB–MreB contacts, with possible secondary effects on the smaller lateral interface |
| **V114E** | **Lateral proto-filament interface** | IA | V118 | None | T116, A117, V118(+), E119, R121, A122(+) | largest lateral MreB–MreB interface, lateral protofilament association |

* Corresponding residue in *C. crescentus* MreB

**(C) ATP-binding residues in *G. stearothermophilus* MreB identified from the ATP-bound crystal structure (PDB 7ZPU; Mao et al., 2023).**

mc, interaction mediated exclusively by main-chain atoms.

**ATP adenine-binding residues:**
K212, R294(mc)
**ATP ribose-binding residues:**
G160(mc), D185, E209, K212
**ATP phosphate-binding residues**
T15, A16(mc), N17(mc), E136, G160(mc), G161(mc), G162(mc), T163, G289, L315(mc)

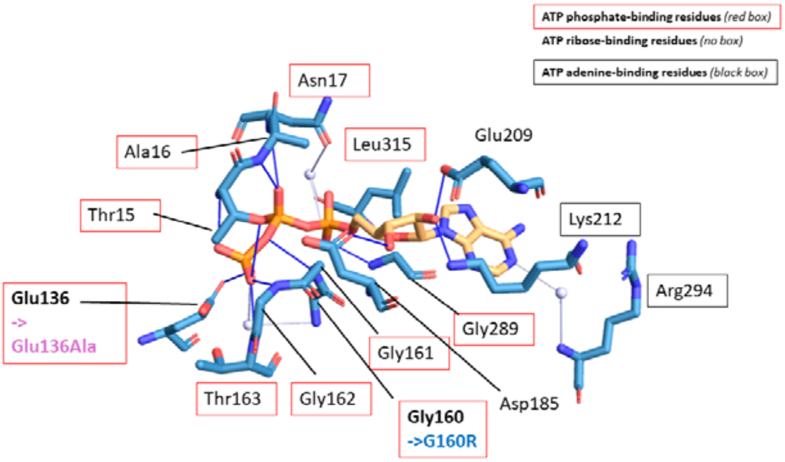


Red and black boxes indicate residues interacting with the ATP phosphates and adenine base, respectively; unboxed residues interact with the ribose.

**(D) Proximity of the analyzed *G. stearothermophilus* MreB ATP-binding-site mutations to nucleotide-interacting residues.** The mutations were mapped relative to residues that directly interact with ATP or AMPPNP, or lie nearby, in ATP-bound *G. stearothermophilus* MreB (PDB 7ZPU) and AMPPNP–$Mg^{2+}$-bound *C. crescentus* MreB (PDB 4CZJ, AMPPNP, a non-hydrolysable ATP analogue).

| **Mutated residue in *Gs*MreB** | | | **Nearby residue(s) (±4 residues) involved in ATP binding in GsMreB-ATP structure (PDB 7ZPU)** | **Corres. In *Cc* *** | **Nearby residue(s) (±4 residues) involved in AMPPNP-$Mg^{2+}$ binding in CcMreB- AMPPNP-$Mg^{2}$ (PDB 4CZJ))** |
|---|---|---|---|---|---|
| **mutation** | **Structural location** | **Sub-domain** | | | |
| G14E | ATP-binding site | IA | Lies near T15, A16(mc), N17(mc), which coordinate ATP β- and γ-phosphates | G18 | Lies near T19, A20(mc), N21, which coordinate AMPPNP a-, β- and γ-phosphates |
| G160R | | IIA | G160(mc) coordinates the ATP ribose O3' and lies near G161(mc), G162(mc), and T163, which coordinate the γ-phosphate and β–γ bridging oxygen | G164 | G164(mc) coordinates waters coordinating ATP γ-phosphate and lies near G165(mc), G166(mc), and T167, which coordinate the γ-phosphate and β–γ bridging oxygen |
| E136A | ATP-binding site | IA | E136 coordinates ATP γ-phosphate | E140 | E140 coordinates waters coordinating AMPPNP γ-phosphate |
| D158A | | IIA | Lies near G160(mc), G161(mc), G162(mc), and T163, which coordinate ATP's ribose O3', γ-phosphate, and β–γ bridging oxygen. | D162 | Lies near G165(mc), G166(mc), which coordinate γ-phosphate, and β–γ bridging oxygen. |
| L171P | Longitudinal proto-fil. interface | IIA | None | L175 | None |
| G231D | | IIB | None | G237 | None |
| V114E | Lateral proto-fil. interface | IA | None | V118 | None |

* Corresponding residue in *C. crescentus* MreB

**REFERENCES:**

Krissinel E, Henrick K. Inference of macromolecular assemblies from crystalline state. J Mol Biol. 2007 Sep 21;372(3):774-97. doi: 10.1016/j.jmb.2007.05.022. Epub 2007 May 13. PMID: 17681537.
